# Late-Onset Preeclampsia is characterised by Accelerated Placental Aging

**DOI:** 10.1101/2025.11.12.687933

**Authors:** Anya L Arthurs, Rudrarup Bhattacharjee, Melanie D Smith, Dulce Medina, Ellen Menkhorst, German Mora, Jessica M Williamson, Lynda K Harris, Jose M Polo, David A MacIntyre, Claire T Roberts

## Abstract

Late-onset preeclampsia (LOPE) is a major pregnancy complication characterised by hypertension and placental dysfunction, resolving only upon delivery. Here, we show that LOPE placentae undergo accelerated molecular aging, marked by telomere attrition, DNA damage and trophoblast senescence. Using primary placental tissue and trophoblast organoids, we demonstrate oxidative stress as a driver of telomere shortening and angiogenic imbalance. Inflammation did not alter placental aging trajectories. Antioxidant treatment (superoxide dismutase) preserved telomere length, reduced DNA damage and restored angiogenic balance, highlighting oxidative stress as a modifiable determinant of placental aging. We identify reduced expression of telomeric repeat-containing RNAs (TERRAs) as a molecular hallmark of LOPE, and show that antisense oligonucleotide-mediated TERRA depletion exacerbates telomere erosion and senescence. Together, these findings delineate oxidative stress and TERRA loss as mechanisms driving placental decline, establish trophoblast organoids as a tractable model of placental aging, and reveal potential therapeutic avenues for mitigating preeclampsia-associated placental dysfunction.

## Introduction

Aging is a fundamental process that drives cellular degeneration, organ dysfunction and systemic health deterioration^1^. The placenta follows a unique aging trajectory whereby as a transient fetal organ, it compresses decades of biological aging into a single gestation. While some deterioration is expected toward term^2^, maternal health^3–8^ and environmental exposures^5,9^ have been shown to hasten placental decline and impair its function. This is characterised by reduced oxygen and nutrient exchange^10^, which contributes to adverse pregnancy outcomes like preeclampsia^11^, fetal growth restriction^12^, preterm birth^13^ and stillbirth^2,14^. These complications reverberate beyond pregnancy, shaping future maternal and offspring health^15–19^, longevity^20^ and social equity^21–24^. With rising maternal age and the global burden of pregnancy disorders^22^, understanding placental aging is ciritical.

Late-onset preeclampsia (LOPE), which arises after 34 weeks of gestation^25^ and accounts for most preeclampsia cases^26,27^, is usually only resolved by delivery of the placenta. Clinically, LOPE is characterised clinically by hypertension and organ dysfunction^25^, while biologically, by oxidative stress^28^, inflammation^29^ and altered trophoblast differentiation^30^. Although telomere shortening^31^ and villous calcification^32^ are accentuated in LOPE compared to gestationally matched controls, the molecular mechanisms that drive accelerated placental aging remain poorly defined. Physiological placental aging normally accelerates after 37 weeks of gestation and coincides with increasing stillbirth risk^2,33^, yet it remains unclear how LOPE diverges from normal aging trajectories at the cellular and molecular level.

Here, we describe accelerated placental aging as a hallmark of LOPE, marked by telomere attrition, cell senescence, DNA damage and altered trophoblast composition. Using trophoblast organoids, we show that oxidative stress is the principal driver of telomere shortening and cellular senescence in LOPE, whereas inflammasome-mediated inflammation does not alter the aging trajectory. Antioxidant treatment preserves telomere length, reduces oxidative stress and ameliorates angiogenic imbalance, highlighting oxidative stress as a tractable therapeutic target. We further identify reduced expression of telomeric repeat-containing RNAs (TERRAs) as an additional hallmark of accelerated placental aging, implicating TERRA loss alongside oxidative stress in driving LOPE pathology. Together, these findings expose unrecognised mechanisms of placental aging and suggest therapeutic avenues for mitigating placental dysfunction in preeclampsia.

## Results

### Accelerated placental aging in LOPE

To investigate whether LOPE is associated with accelerated placental aging, we analysed term placentae, both from healthy (control) and LOPE pregnancies collected across 37– 42 weeks’ gestation. Telomere length assessment showed a gestational age–dependent decline from 39 weeks’ gestation onwards in all placentae, with LOPE samples exhibiting consistently shorter telomeres than controls (mean 3.68 kB versus 4.12 kB across all gestational ages, respectively; **Fig. 1a**). Cellular senescence, assessed by SA-β-gal staining, was significantly higher in LOPE placentae, with mean 43.2% of trophoblasts positive compared to mean 31.8% in controls (**Fig. 1b**).

**Figure 1:**
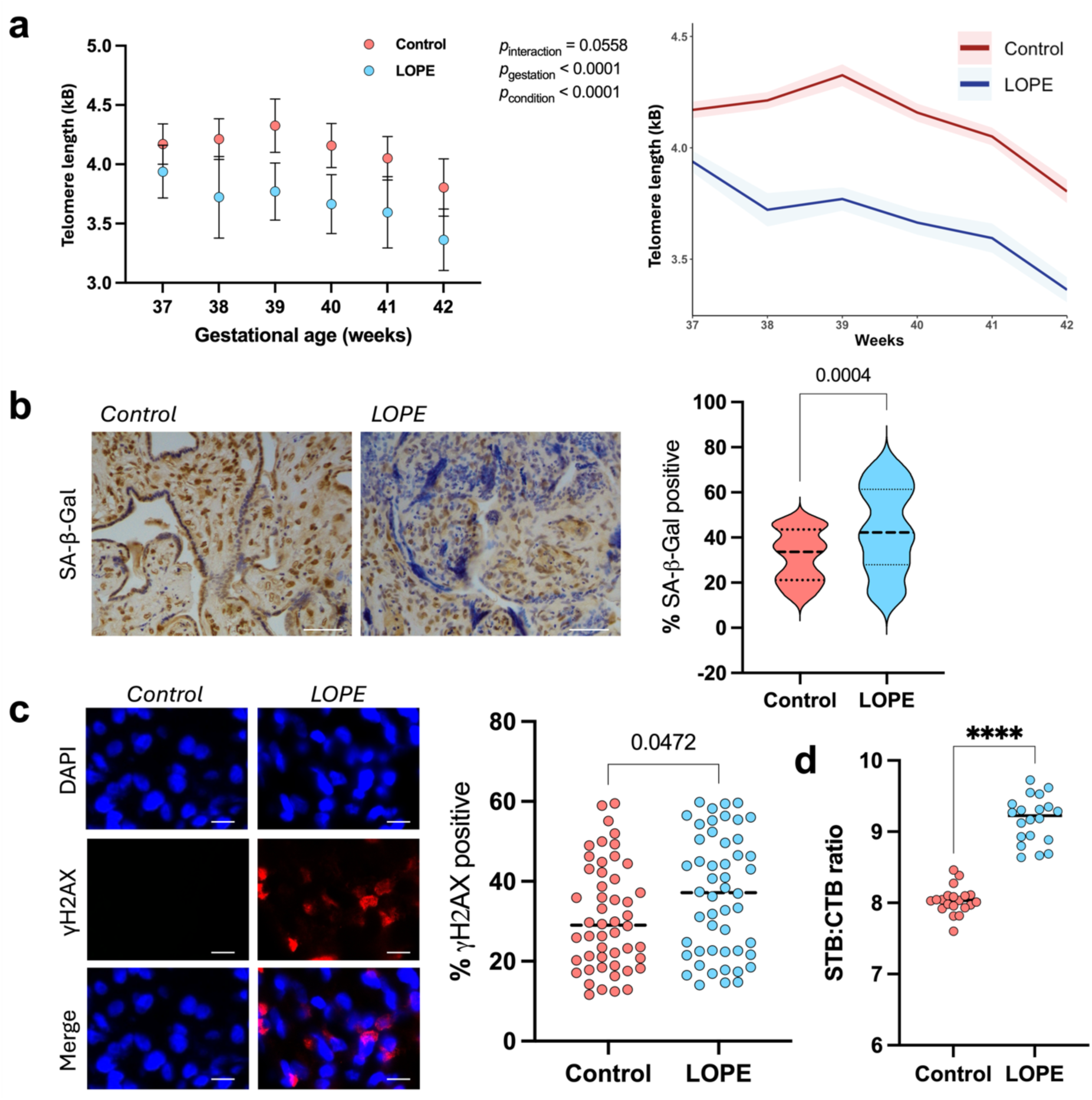
LOPE placentae display telomere attrition, increased senescence, and altered trophoblast composition at term. In placentae delivered at 37 weeks’ gestation from control (red) and LOPE (blue) pregnancies: **a**, mean telomere length (kB), assessed using a two-way ANOVA. **b**, SA-β-Gal (violet) counterstaining on pan cytokeratin (brown) determined senescence. Scale bar, 150 µm **c**, γH2AX (red) immunofluorescence determined double stranded DNA breaks. Representative images *[left]* scale bar, 10 µm; *[right]* quantified data. **d**, Immunohistochemical analysis determined the ratio of syncytiotrophoblasts to cytotrophoblast cells, **** *p* < 0.0001. For **b-d**, assessed using Mann-Whitney tests. For **b-c**, n=10 fields of view were selected for each sample and mean positive staining was recorded. For **d**, n=10 fields of view were selected for each sample and all nuclei were counted within a field of view. Data in **a-b** are expressed in violin plots, **c-d** are expressed in scatter plots. n=50/condition for all. Source data file is provided.

Elevated levels of DNA damage were confirmed through increased γH2AX foci, a marker where H2AX is phosphorylated at Ser139 in response to DNA damage. LOPE placentae had a higher proportion of γH2AX-positive nuclei (median 37.1%) compared to gestational age–matched controls (median 29.1%; **Fig. 1c**). To evaluate trophoblast differentiation, we calculated the syncytiotrophoblast-to-cytotrophoblast (STB:CTB) ratio, which normally increases toward term as cells undergo terminal differentiation. LOPE placentae showed a higher STB:CTB ratio (median 9.27) compared to controls (median 8.03; **Fig. 1d**), consistent with accelerated trophoblast maturation. DNA methylation levels, another recognised hallmark of cell aging^34^, was also assessed in control and LOPE placentae. However, no statistically significant changes in methylation were observed (**Supplementary Figures 6-G**). This was not unexpected, given conflicting reports of DNA methylation in aging LOPE placentae^35–37^.

Together, these results demonstrate that LOPE placentae exhibit premature and accelerated aging at term, marked by telomere attrition, increased cellular senescence and DNA damage, as well as altered trophoblast composition.

### Establishment of an *in vitro* placental aging model

We first sought to validate trophoblast organoids derived from primary placental tissue as an *in vitro* model of *in vivo* placental aging. Trophoblast organoids were generated from both healthy and LOPE placenta tissues collected from deliveries at 37 weeks’ gestation. Organoids were cultured for 4 weeks with a subset of organoids harvested each week for DNA extraction (**Supplementary Figure 2c**). Weekly telomere length measurements confirmed that trophoblast organoids mirrored the progressive telomere attrition observed *in vivo,* with telomeres in LOPE-derived trophoblast organoids consistently shorter (mean telomere length at 4 weeks in culture: 3.54 kB) compared to controls (mean telomere length at week 4: 3.81 kB; **Supplementary Figure 2a,b**).

Notably, there was no significant difference in telomere length between trophoblast organoids derived from 37-weeks deliveries and cultured for 4 weeks, and trophoblast stem cells collected directly from 40-weeks births, confirming that the *in vitro* system faithfully recapitulates the *in vivo* aging trajectory.

### Antioxidant treatment rescues telomere attrition and senescence in LOPE trophoblast organoids

Oxidative stress is widely implicated as a primary driver of LOPE pathology^38^. To investigate whether oxidative stress drives accelerated placental aging, we generated trophoblast organoids from control and LOPE placentae delivered at 37 weeks’ gestation, and treated them with the antioxidant superoxide dismutase (SOD) throughout the 4 week culture period (**Fig. 2a,b**). SOD catalyses the conversion of superoxide radicals to oxygen and hydrogen peroxide.

**Figure 2:**
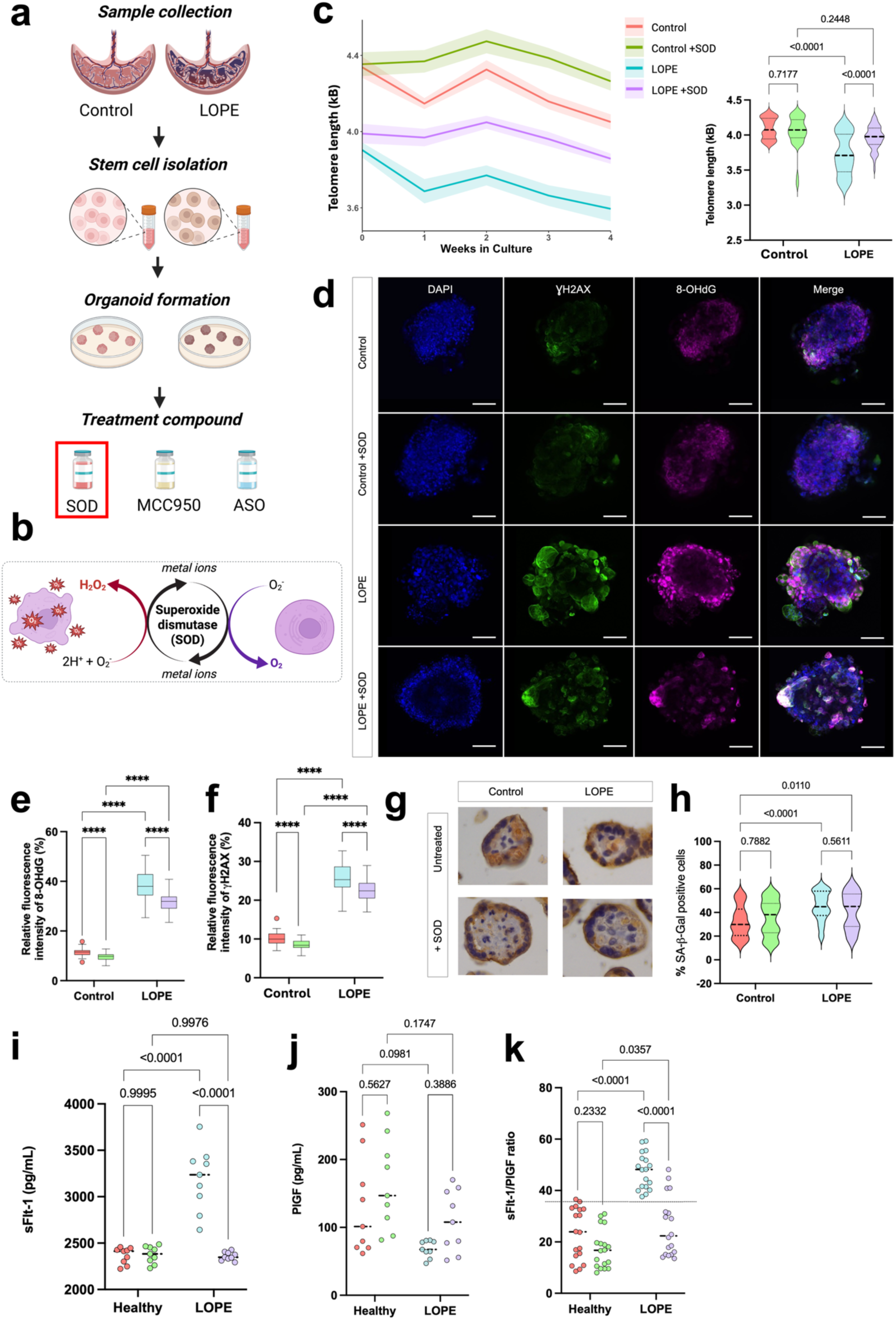
Antioxidant treatment with superoxide dismutase (SOD) preserves telomere length, reduces oxidative stress and senescence, and improves angiogenic balance in LOPE trophoblast organoids. **a**, Workflow for sample collection, trophoblast stem cell isolation, organoid formation and antioxidant treatment with superoxide dismutase (SOD). **b**, Schematic of SOD catalysing the dismutation of superoxide anions into oxygen. **c**, Telomere length dynamics over 4 weeks in culture (left) and violin plots of telomere length at endpoint (right) in control and LOPE organoids with or without SOD treatment. **d**, Representative immunofluorescence images of control and LOPE trophoblast organoids stained for γH2AX (DNA damage, green), 8-OHdG (oxidative stress, magenta) and DAPI (nuclei, blue) in the presence or absence of SOD. Scale bar, 100 µm. **e,** Quantification of 8-OHdG and **f,** γH2AX fluorescence intensity in control and LOPE trophoblast organoids with or without SOD treatment. Data presented as Tukey’s box and whisker plots. **g**, Representative immunohistochemistry images showing SA-β-Gal (violet) to assess senescence, co-stained with pan-cytokeratin to image trophoblast cells and counterstained with DAPI for nuclei; imaged at 40× magnification, and **h,** quantification of SA-β-Gal-positive staining. **i-k,** Concentrations of angiogenic factors in culture media after 4 weeks, including **i,** sFlt-1 (pg/mL), **j,** PlGF (pg/mL), and **k,** sFlt-1/PlGF ratio, in control and LOPE trophoblast organoids with or without SOD treatment. Data are presented as mean ± s.d. Trophoblast organoids were generated from n = 3 separate biological samples per fetal sex, per group; experiments performed in technical triplicate. For **c**,**e**-**f**,**h**,**i-k,** statistical analysis was performed using Welch’s ANOVA with multiple comparisons. \*\*\*\**p* < 0.0001; ns, not significant. Source data file is provided.

Across four weeks in culture, LOPE-derived trophoblast organoids displayed significantly shorter telomeres compared to controls (mean 3.21 kb versus 3.84 kb at week 4). After 4 weeks in culture, telomere length in LOPE trophoblast organoids was preserved with SOD (mean 3.62 kb with SOD versus 3.94 kb without; **Fig. 2c**), partially restoring LOPE values to control levels.

We next assessed oxidative stress and DNA damage. Immunofluorescence revealed significantly increased γH2AX and 8-OHdG staining in LOPE trophoblast organoids compared to controls, consistent with elevated DNA damage and oxidative stress. These markers were markedly reduced with SOD treatment in both groups (**Fig. 2d**). Quantitative analysis confirmed significant decreases in 8-OHdG intensity (−17.6%) and γH2AX intensity (−13.2%) following antioxidant treatment in LOPE organoids (**Fig. 2e,f**).

Cellular senescence, assessed by SA-β-Gal staining, was significantly higher in LOPE trophoblast organoids compared to controls (median 61.5% versus 39.8% positive cells). However, SOD treatment did not significantly alter the amount of SA-β-Gal staining for either group (**Fig. 2g,h**).

We next examined angiogenic and anti-angiogenic factors in organoid culture media as secreted by trophoblast organoids.This is because the soluble fms-like tyrosine kinase 1 (sFlt-1) / placental growth factor (PlGF) ratio in maternal circulation is the most widely used clinical biomarker for preeclampsia and reflects placental dysfunction^39^. A clinical cut-off of 38 or higher signifies elevated risk of preeclampsia (indicated in diagrams with a dotted line, e.g. **Fig. 2k**). Furthermore, excess placental sFlt-1 exacerbates features of preeclampsia^40^.

sFlt-1 concentration was elevated in culture media of organoids from LOPE placentae compared with healthy controls, whereas PlGF was unchanged, resulting in an increased sFlt-1/PlGF ratio (**Fig. 2i–k**). These changes were partially ameliorated following SOD treatment of LOPE trophoblast organoids, linking oxidative stress and telomere attrition with dysregulated release of angiogenic factors.

Taken together, these findings demonstrate that oxidative stress is one of the key drivers of telomere attrition, DNA damage, senescence, and secretory dysfunction in LOPE trophoblast organoids. Antioxidant treatment with SOD conferred significant protection and halted aging phenotypes.

### NLRP3 inflammasome inhibition does not rescue telomere attrition nor senescence in LOPE trophoblast organoids

We next tested whether targeting inflammatory pathways could alter the trajectory of accelerated placental aging in LOPE. The NLRP3 inflammasome has been implicated in LOPE pathogenesis^41–43^. To test whether inflammatory pathways contribute to accelerated placental aging, we treated both control and LOPE trophoblast organoids with the selective NLRP3 inflammasome inhibitor MCC950 (**Fig. 3a,b**). Immunoblotting confirmed that MCC950 reduced NLRP3 protein levels in treated organoids (**Fig. 3e**), validating effective inflammasome inhibition.

**Figure 3:**
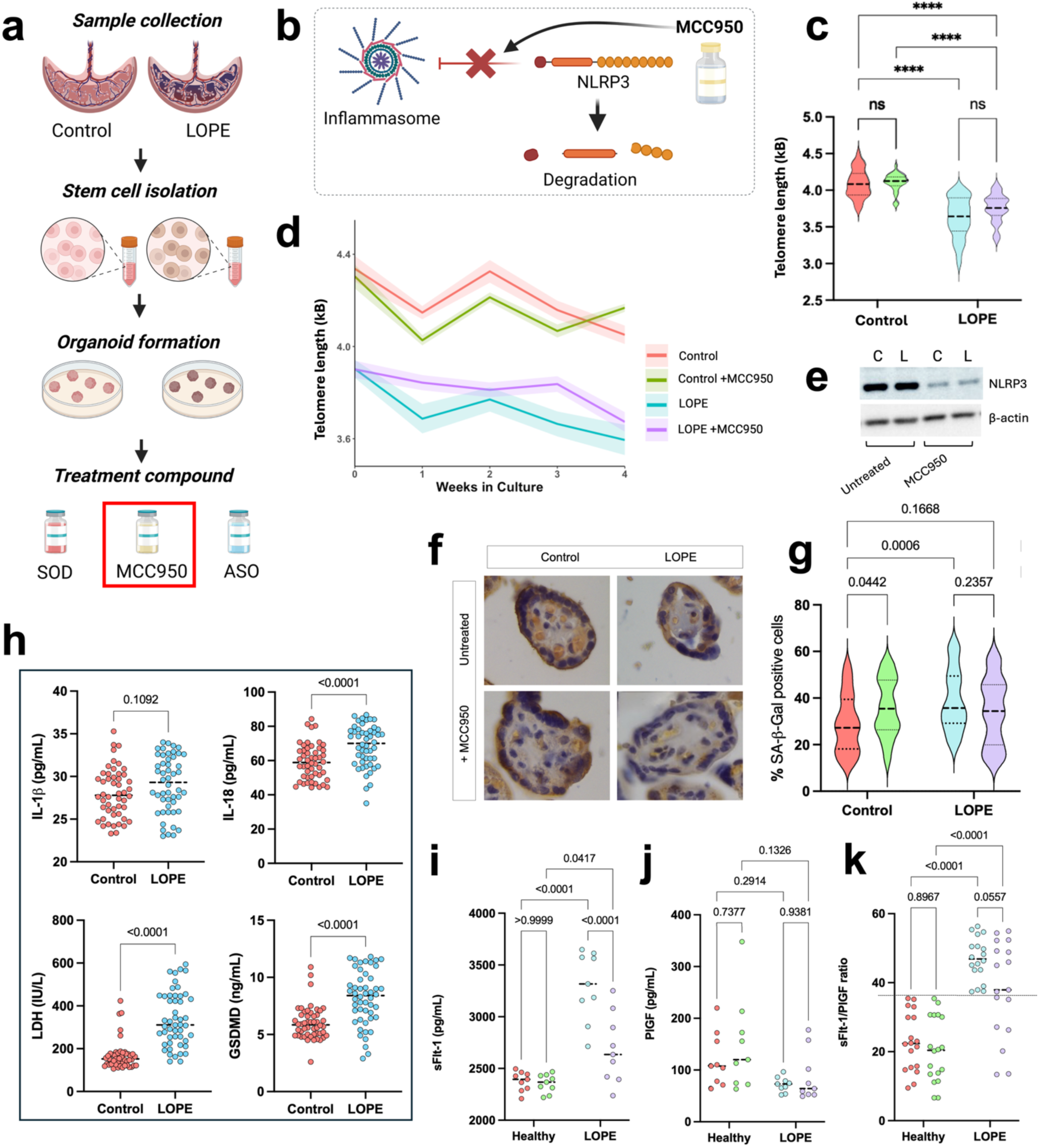
Inhibition of the NLRP3 inflammasome with MCC950 reduces NLRP3 abundance, but does not rescue LOPE trophoblast organoids from accelerated aging or angiogenic imbalance. **a**, Workflow for sample collection, trophoblast stem cell isolation, organoid formation and treatment with MCC950. **b**, Schematic of MCC950 blocking inflammasome assembly and promoting NLRP3 degradation. **c**, Violin plots of telomere length at endpoint and **d,** telomere length dynamics over 4 weeks in culture in control and LOPE organoids with or without MCC950 treatment. **e**, Representative Western blot showing NLRP3 protein levels in control (C) and LOPE (L) trophoblast organoids with or without MCC950 treatment. Molecular weight of NLRP3 ∼119 kDa, β-actin ∼42 kDa. **f**, Representative immunohistochemistry images showing SA-β-Gal (violet) counterstaining on pan cytokeratin (brown) to assess senescence, imaged at 40× magnification and **g,** quantification of SA-β-Gal-positive staining (violin plot). **h,** Scatter plots depicting concentrations of maternal circulating inflammatory markers associated with NLRP3 inflammasome activation, including IL-1β (top left), IL-18 (top right), lactate dehydrogenase (LDH, bottom left) and GSDMD (bottom right), in healthy (control) and LOPE pregnancies. **i-k**, Scatter plots depicting concentrations of angiogenic factors in culture media after 4 weeks, including **i**, sFlt-1, **j,** PlGF and **k,** sFlt-1/PlGF ratio, in control and LOPE trophoblast organoids with or without MCC950. Trophoblast organoids were generated from n = 3 separate biological samples per fetal sex, per group; experiments performed in technical triplicate. For **c**,**g**,**i-k,** statistical analysis was performed using Welch’s ANOVA with multiple comparisons. For **h,** n=50 per group, data assessed using Mann-Whitney tests. \*\*\*\**p* < 0.0001; ns = not significant. Source data file is provided.

Across four weeks in culture, LOPE-derived trophoblast organoids exhibited significantly shorter telomeres compared to controls (week 4 mean 3.58 kb versus 4.09 kb). MCC950 treatment did not preserve telomere length in either group (week 4 mean telomere length: 4.14 kb versus 4.09 kb in controls, 3.60 kb versus 3.58 kb in LOPE; **Fig. 3c,d**). Despite the reduced NLRP3 levels, attrition of telomeres persisted suggesting that NLRP3-mediated inflammation is not a primary driver of this aging phenotype.

Cellular senescence, measured by SA-β-Gal staining, was higher in LOPE trophoblast organoids compared to controls (median 38.2% versus 26.5% positive cells). MCC950 treatment did not alter senescent cell burden in LOPE trophoblast organoids, and increased cellular senescence in control organoids (36.1% with MCC950 treatment versus 26.5% in untreated) (**Fig. 3f,g**). MCC950 did not reduce DNA damage nor oxidative stress, as both γH2AX- and 8-OHdG–positive nuclei remained unchanged relative to untreated trophoblast organoids (**Supplementary Fig. 4**).

Given the established role of inflammasome activation in LOPE pathogenesis, we next assessed maternal circulating mediators of inflammasome activity. Levels of interleukin 1β (IL-1β), interleukin 18 (IL-18), gasdermin D (GSDMD) and lactate dehydrogenase (LDH) were measured in whole blood samples collected from pregnant women at 34 weeks’ gestation. IL-18, GSDMD and LDH were significantly elevated in the maternal blood from LOPE pregnancies compared with healthy controls, whereas IL-1β remained unchanged (**Fig. 3h**). These results indicate that maternal inflammation is present *in vivo*, but that direct inhibition of the inflammasome *in vitro* is insufficient to rescue accelerated aging phenotypes.

Similar to the above antioxidant study, we evaluated whether inflammasome inhibition modulated the sFlt-1/PlGF ratio (as a preeclampsia biomarker). As expected, sFlt-1 was elevated and PlGF reduced in culture media of organoids derived from placentae of LOPE pregnancies, resulting in an increased sFlt-1/PlGF ratio. MCC950 treatment did not significantly change sFlt-1, PlGF, nor their ratio in either control or LOPE trophoblast organoids (**Fig. 3i-k**).

Taken together, these results show that although MCC950 effectively inhibits the NLRP3 inflammasome, it does not prevent telomere shortening, senescence, nor dysregulated angiogenic factor release in LOPE trophoblast organoids. Combined with earlier results, this further highlights that oxidative stress, rather than inflammasome-mediated inflammation, is a dominant driver of accelerated placental aging.

### SOD treatment promotes syncytial knot formation in LOPE trophoblast organoids

Syncytial knots, aggregates of condensed syncytiotrophoblast nuclei shed from villus surfaces, are a histological hallmark of accelerated placental aging^44^. To determine whether oxidative stress contributes to this process, we examined syncytial knot formation in control and LOPE trophoblast organoids treated with either MCC950 or SOD (**Fig. 4a**).

**Figure 4.**
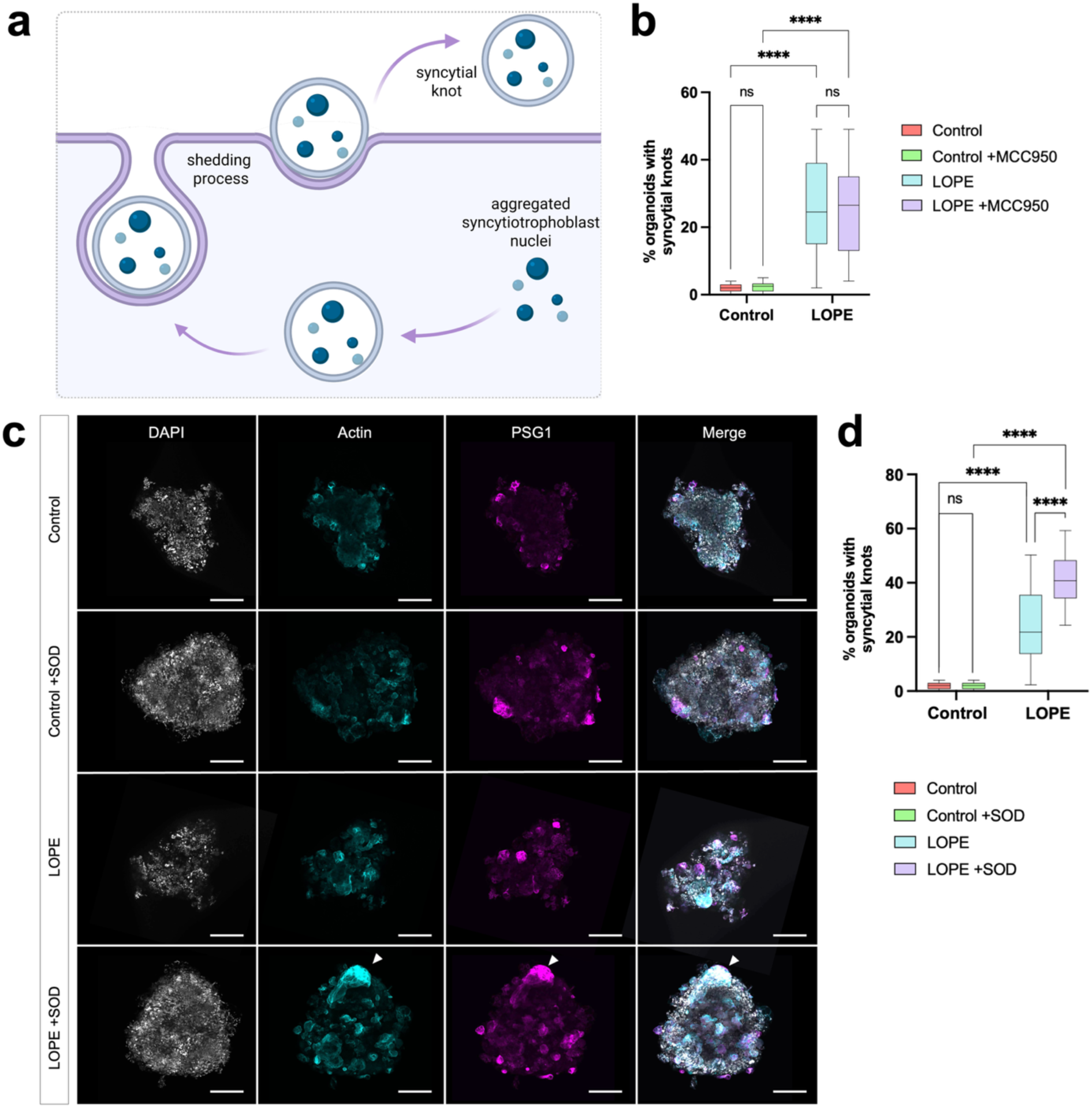
**Antioxidant treatment with SOD, but not MCC950, increases syncytial knot formation in LOPE trophoblast organoids**. **a**, Schematic depicting syncytial knot formation and shedding. **b,d,** Quantification of syncytial knot density in control and LOPE organoids with or without treatment with **b,** MCC950 and **d,** SOD. Data presented as Tukey’s box and whisker plots. **c,** Representative confocal images of trophoblast organoids stained with DAPI (nuclei, white), actin (cyan) and PSG1 (syncytiotrophoblast, magenta). Merge images show syncytial knot formation, indicated by white arrowheads. Data are presented as mean ± s.d. (n = 50 organoids per group, quantified blinded to condition) and compared using Welch’s ANOVA with multiple comparisons. ****p < 0.0001; ns, not significant. Scale bar, 100 µm. Source data file is provided.

#### MCC950 does not alter syncytial knot frequency

Quantification revealed that syncytial knots were rarely detected in control trophoblast organoids and remained infrequent following MCC950 treatment. However, LOPE organoids displayed a significantly higher proportion of syncytial knots, consistent with their aged phenotype, and this was not reduced by MCC950 (**Fig. 4b**). These findings indicate that inflammasome inhibition does not alter syncytial knot formation frequency.

#### SOD exacerbates syncytial knot burden

We next assessed whether antioxidant treatment could modulate syncytial knot abundance. Confocal imaging of trophoblast organoids stained for DAPI, actin and PSG1 revealed characteristic nuclear aggregates consistent with syncytial knots in LOPE organoids, which were notably increased following SOD treatment (**Fig. 4c**). Quantitative analysis confirmed that SOD significantly increased the proportion of organoids with syncytial knots in LOPE, while no effect was observed in controls (**Fig. 4d**).

Taken together, these results demonstrate that SOD treatment promotes syncytial knot formation in LOPE trophoblast organoids, and that this phenotype is not affected by inflammasome inhibition.

### Loss of TERRA contributes to telomere attrition and senescence in LOPE trophoblast organoids

Telomeric repeat–containing RNAs (TERRAs) form R-loops at chromosome ends and stabilise telomeres. Given the altered telomere length observed in primary LOPE and healthy (control) placenta samples, we assessed TERRA expression in these samples. qPCR analysis showed that TERRAs were significantly downregulated in primary LOPE placenta tissue compared with healthy controls. Consistently, we observed reduced expression of 20q, 10q, 13q and XY TERRAs in LOPE trophoblast organoids compared to controls (**Fig. 5a–d**).

**Figure 5.**
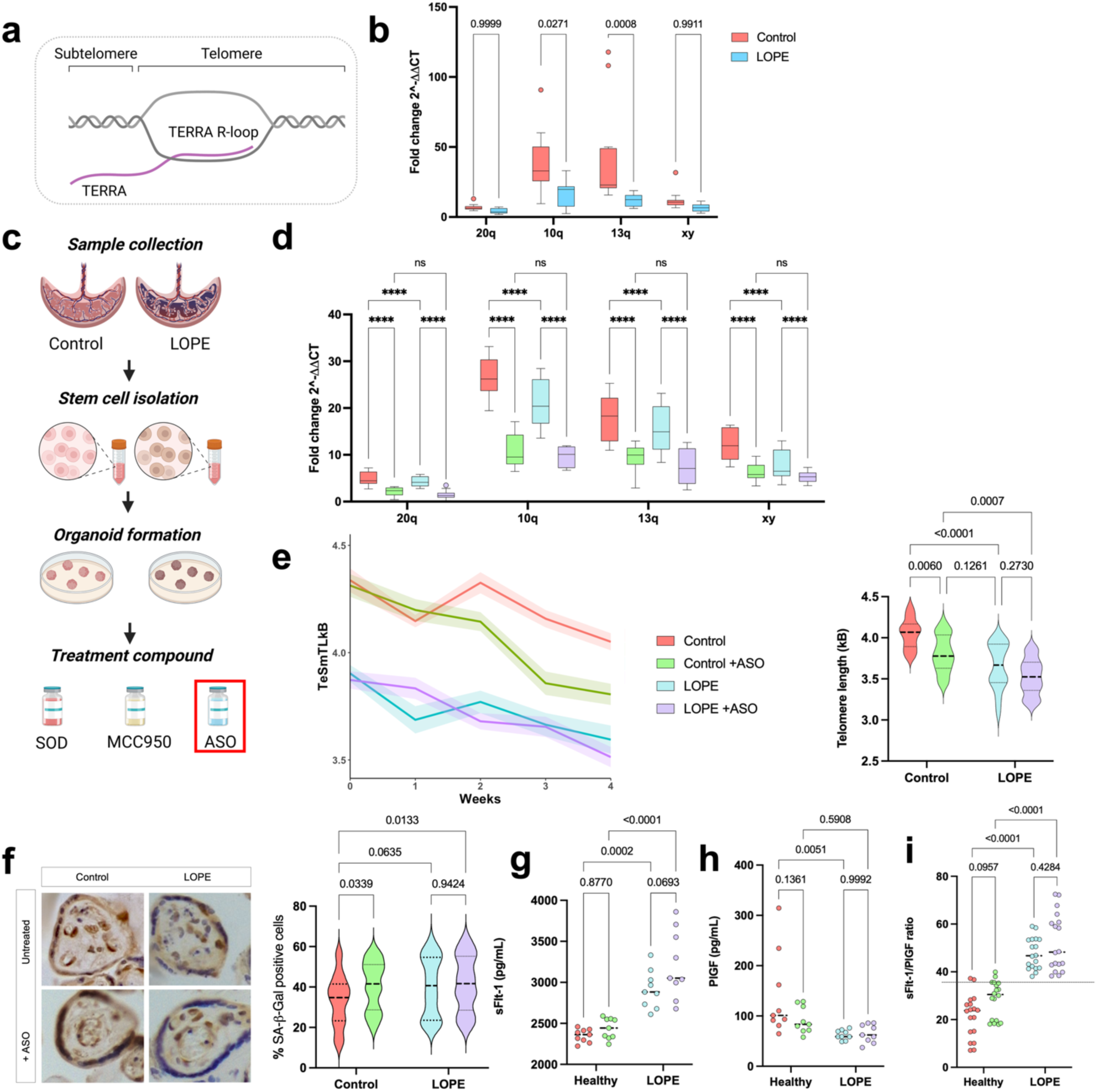
Antisense oligonucleotide (ASO)–mediated inhibition of TERRAs reduces telomere length and increases senescence and angiogenic imbalance in LOPE trophoblast organoids. a, Schematic depicting TERRA R-loop formation at the telomere. **b,** Expression of 20q, 10q, 13q and XY TERRAs in primary placental tissue from control and LOPE pregnancies, assessed using qPCR. **c,** Workflow for sample collection, trophoblast stem cell isolation, organoid formation and treatment with antisense oligonucleotide (ASO). **d,** Expression of 20q, 10q, 13q and XY TERRAs in trophoblast organoids from control and LOPE pregnancies with or without ASO treatment, assessed using qPCR. **e,** Telomere length dynamics over 4 weeks in culture (left) and violin plots of telomere length at endpoint (right) in control and LOPE organoids with or without ASO treatment. **f,** Representative immunohistochemistry images showing SA-β-Gal (violet) counterstaining on pan cytokeratin (brown) to assess senescence, imaged at 40× magnification (left), and quantification of SA-β-Gal–positive staining (right). **g–i,** Concentrations of angiogenic factors in culture media after 4 weeks, including **g,** sFlt-1 **h,** PlGF and **i,** sFlt-1/PlGF ratio, in control and LOPE trophoblast organoids with or without ASO treatment. Data are presented as mean ± s.d. Trophoblast organoids were generated from n = 3 separate biological samples per fetal sex, per group; experiments performed in technical triplicate. Primary tissue samples, n=50 per group. For **b, d,** data are presented as Tukey’s box and whisker plots. For **b,d–i,** statistical analysis was performed using Welch’s ANOVA with multiple comparisons. \*\*\*\**p* < 0.0001; ns, not significant. Source data file is provided.

To test whether further depletion of TERRA would influence aging trajectories, we treated control and LOPE organoids with antisense oligonucleotides (ASOs) targeting TERRAs (**Fig. 5c**). After four weeks in culture, ASO treatment exacerbated telomere shortening in both control and LOPE organoids (**Fig. 5d**), consistent with the role for TERRAs in maintaining telomere length.

We next examined whether TERRA depletion influenced cellular senescence. SA-β-Gal staining revealed an increased proportion of senescent cells in LOPE organoids compared with controls, which was further augmented following ASO treatment (**Fig. 5f**). These results suggest that TERRA loss contributes to premature trophoblast senescence in LOPE. TERRA depletion via ASO also did not reduce DNA damage or oxidative stress, as γH2AX- or 8-OHdG–positive nuclei remained unchanged relative to untreated trophoblast organoids (**Supplementary Fig. 4**).

As angiogenic dysfunction is a hallmark of LOPE, we assessed whether TERRA depletion altered release of placental factors. In culture media, sFlt-1 was elevated, PlGF was reduced, and the sFlt-1/PlGF ratio was significantly increased in LOPE pregnancies compared with healthy controls (**Fig. 5g–i**). ASO treatment of control organoids partially recapitulated these changes, with higher sFlt-1 and reduced PlGF secretion, leading to an increased ratio.

Taken together, these findings demonstrate that TERRA loss contributes to telomere attrition, senescence and dysregulated angiogenic signalling in LOPE trophoblast organoids, establishing TERRA as an important regulator of placental aging.

## Discussion

Here we demonstrate that LOPE is characterised by accelerated placental aging, marked by telomere attrition, increased senescence, DNA damage and altered trophoblast composition at term. Using primary placentae and trophoblast organoids, we show that oxidative stress and loss of TERRA downregulation, rather than inflammasome-mediated inflammation, are the dominant drivers of these aging phenotypes. These findings provide direct mechanistic evidence distinguishing LOPE from physiological placental aging and identify new molecular signatures of placental decline.

Our data indicate that oxidative stress accelerates trophoblast senescence through telomere attrition, consistent with studies in other tissues where oxidative stress promotes cellular aging and telomere erosion. In parallel, we observed reduced expression of telomeric repeat-containing RNAs (TERRAs) in LOPE, which may represent an additional hallmark of accelerated placental aging. While the relationship between oxidative stress and TERRA loss remains unresolved, these changes together suggest multiple, converging pathways of telomere dysfunction in LOPE. In contrast, inhibition of the NLRP3 inflammasome did not prevent telomere shortening, senescence nor angiogenic imbalance in LOPE organoids. NLRP3 activation in placental tissue induces the release of pregnancy-compromising factors, including sFlt-1, IL-1β and other pro-inflammatory cytokines and alarmins, into the maternal circulation, driving oxidative stress, systemic inflammation and endothelial dysfunction^45,46^. We observed this systemic inflammatory signature in maternal blood. However, once placental tissue was removed from maternal circulation and cultured as organoids, direct inflammasome inhibition no longer altered the aging trajectory. This divergence may reflect the complexity of the *in vivo* environment. Collectively, our findings suggest that while maternal systemic inflammation amplifies placental decline *in vivo*, oxidative stress within trophoblasts remains the proximate driver of accelerated aging.

The potential therapeutic implications of these findings are underscored by our antioxidant rescue experiments. Treatment with SOD preserved telomere length, reduced oxidative DNA damage, alleviated trophoblast senescence and partially normalised angiogenic factor secretion in LOPE organoids. Unexpectedly, however, syncytial knot formation increased following SOD treatment, despite improvements in other aging parameters. This contrasts with the broadly protective effects of antioxidant therapy but suggests that knot formation may not solely represent a pathological endpoint. Instead, syncytial knots may function adaptively by clearing nuclear debris and maintaining syncytiotrophoblast homeostasis under conditions of stress. The observation that knots also accumulate in normal human placentae at term^44^ supports the interpretation that their formation in SOD-treated organoids reflects physiological tissue turnover rather than accelerated degeneration. Furthermore, as SOD does not increase knots in controls, its therapeutic potential still remains very high as it would not otherwise negatively impact placental function. Together, these findings indicate that antioxidant therapy can mitigate telomere dysfunction and senescence while simultaneously activating adaptive clearance mechanisms that may contribute to placental resilience.

The clinical relevance of these findings lies in the potential to target oxidative stress as a modifiable driver of placental aging. Although antioxidant therapies have shown mixed results in pregnancy, our data suggest that selective strategies, timed to the late pregancy window when LOPE emerges, may hold greater promise. Integration of biomarkers such as the sFlt-1/PlGF ratio could enable risk stratification and identify women most likely to benefit from intervention. By refining therapeutic timing and patient selection, interventions that mitigate placental oxidative stress may reduce adverse outcomes associated with LOPE and provide broader insights into managing aging-related placental dysfunction.

In addition, our observation of reduced TERRA expression identifies an RNA-based therapeutic opportunity. While global TERRA overexpression is undesirable, as excessive TERRA levels are associated with cancer^47^, a more targeted approach could protect endogenous TERRAs from degradation. We propose that steric hindrance antisense oligonucleotides (ASOs) could be designed to bind TERRAs at promoter regions, shielding them from endogenous decay pathways such as nonsense-mediated decay without altering their normal function. Such an approach could selectively stabilise TERRA transcripts in placenta, preserve telomere integrity and delay placental senescence, providing a novel therapeutic avenue distinct from direct overexpression strategies.

Several limitations should be acknowledged. Our analyses were performed at delivery and in organoid models, which may not fully capture earlier gestational dynamics of placental aging. While oxidative stress emerged as the dominant driver, additional maternal and fetal influences may contribute to disease progression *in vivo*. Moreover, the precise mechanisms by which TERRA loss accelerates telomere dysfunction remain to be elucidated, and the feasibility of ASO-based interventions requires extensive preclinical validation. Future studies incorporating longitudinal sampling across gestation and functional interrogation of TERRA biology will be critical to delineate the trajectory and regulation of LOPE-associated aging.

In summary, this study identifies oxidative stress and TERRA reduction as central accelerants of placental aging in LOPE, distinguishing pathological from physiological trajectories of decline. By establishing trophoblast organoids as a tractable model, we reveal oxidative stress as a modifiable target and propose TERRA-stabilising ASOs as a potential therapeutic strategy. Moreover, our data highlight that syncytial knot formation may represent an adaptive clearance mechanism rather than a purely degenerative hallmark, offering new perspectives on how the placenta maintains tissue homeostasis under stress. These findings expand our understanding of reproductive aging and suggest new opportunities to mitigate placental dysfunction and its consequences for maternal and fetal health.

## Methods

### Human participants and samples

Term placentae and maternal blood samples were obtained with informed consent after delivery from the Lyell McEwin Hospital, in Elizabeth, South Australia and from Flinders Medical Centre, in Bedford Park, South Australia. Ethics were obtained from the Women’s and Children’s Health Network Human Research Ethics Committee (HREC/14/WCHN/90) and the Southern Adelaide Local Health Network Human Research Ethics Committee (2024/HRE00218), respectively.

LOPE was defined as the onset of preeclampsia after 34 weeks’ gestation. Preeclampsia was defined as (peripheral) hypertension [systolic BP (SBP) ≥ 140 mmHg or diastolic BP (DBP) ≥ 90 mmHg] after 20 weeks of gestation in previously normotensive women, with proteinuria (24 h urinary protein ≥ 300 mg or spot urine protein: creatinine ratio ≥ 30 mg/mmol creatinine or urine dipstick protein ≥ 2+) or any multi-organ complication of preeclampsia^48^. The estimated date of delivery was calculated from a certain last menstrual period (LMP) date and was only adjusted if either (1) a scan performed at < 16 weeks of gestation found a difference of ≥7 days between the scan gestation and that calculated by the LMP or (2) on 20-week scan a difference of ≥ 10 days was found between the scan gestation and that calculated from the LMP. If the LMP date was uncertain, then scan dates were used to calculate the estimated date of delivery. The clinical characteristics of all the participants are summarized in **Supplementary Table 1**.

Term placental tissues were collected within 1-6 hours of delivery, where a section of villous tissue was biopsied (∼0.1g). Samples were then washed in Dulbecco’s phosphate-buffered saline (DPBS) and transported to the laboratory on ice. Term placental tissues that were not used to generate organoids were snap frozen in liquid nitrogen for 15 mins prior to storage at -80°C.

Maternal blood samples, collected from participants at ∼34 weeks’ gestation, were immediately transported to the laboratory on ice. Whole blood samples were aliquoted into vials and stored at -80°C.

### Derivation and culture of trophoblast organoids from human placental samples

Trophoblast organoids were generated from full term (37 weeks’ and 40 weeks’ gestation) healthy and preeclamptic placentae as per Yang, *et al*.^49^ with alterations to culture conditions. Briefly, placental villi were dissected and washed in HAMS/F12 (#11765054, Gibco) prior to dissociation using a 0.2% Trypsin / 0.02% EDTA solution. After filtration, undigested tissue was further processed using collagenase V. Pooled isolated cells were centrifuged (400 xg, 5 min) and washed (Advanced DMEM/F12 medium, #12634010, Gibco).

Trophoblast stem cells were then seeded at 1×10^5^ cells per well with 3 mL total organoid media (TOM as per Yang, *et al*. with suggested media additions^49^) in a Corning^®^ Costar^®^ Ultra-Low Attachment Multiple Well Plate (#CLS3471, Merck) and placed on an orbital shaker (85 rpm, Thermo Scientific™, Cat. 88881102, orbit 1.9 cm) in an incubator (37°C, 5% O_2_, 5% CO_2_). Trophoblast stem cells were cultured for 1 week for formation (termed Week 0) prior to treatment/assessment. TO polarity was confirmed using immunofluorescence microscopy (**Supplementary Figure 1**).

### Compound treatments

Trophoblast organoids were treated with one of three compounds:

1. MCC950 (1 μM; Selleck Chemicals, USA, #S7809). MCC950 targets NLRP3, inhibiting NLRP3 inflammasome assembly and resulting in NLRP3 degradation. Dosage was as per Perera, *et al.*^50^
2. Superoxide dismutase (SOD, 30 U/mL; Active Recombinant Human Superoxide Dismutase #SOD1-8H, Creative Biomart, New York, USA). SOD is an enzyme that alternately catalyses the dismutation of the superoxide anion radical, resulting in antioxidant effects.
3. Antisense oligonucleotides (ASO): TERRA depletion was achieved as previously described^51,52^ with some modifications. 200nM ASO LNA Gapmer Oligos (Qiagen) specifically designed to inhibit TERRAs, and Oligofectamine Transfection Reagent (Thermo Fisher Scientific) were added to culture media and allowed to incubate with trophoblast organoids on the orbital shaker. Scrambled LNA was used as a negative control (results in **Supplementary Figure 4**). Sequences of oligos can be found in **Supplementary Table 2.**

Trophoblast organoids were treated for 4 weeks of culture (termed Week 1-4). Dosages were sourced from established protocols^50,53^. Trophoblast organoids were then harvested for telomere length assessment and immunofluorescence.

### Telomere length assays

DNA was isolated from placenta tissues using the Qiagen DNeasy Blood and Tissue Kit (Cat ID #69504; Qiagen), following the manufacturer’s instructions for animal tissues. DNA concentration was quantified using the Nanodrop 2000 Spectrometer (Thermo Fisher Scientific). DNA quality was monitored by agarose gel electrophoresis. No sample had to be discarded due to DNA degradation.

Telomere length in placental DNA samples was measured as per Arai *et al.*^54^ and Martin *et al*.^55^ For primary placenta tissue, n=50 per gestational week at time of delivery, for control and LOPE; total n=600. For trophoblast organoids, organoids from n=3 patients per fetal sex, per treatment group, performed in technical triplicate; total n=216. Weekly telomere length measurements were performed over Weeks 0-4 of culture, therefore final n=1080. Briefly, telomere length was measured as abundance of telomeric sequence with that of the single-copy reference gene 36B4 using monochrome multiplex qPCR. Primer sequences are listed in **Supplementary Table 2**. Each plate included three internal control DNA samples of known length (6.5 kb, 3.5 kb and 2.0 kb) to correct for plate-to-plate variation. PCR reactions were performed in triplicate on an Applied Biosystems 7900HT Fast Real-Time PCR system (384-well format). The intra-assay and inter-assay coefficients of variation were 2.3% and 4.9%, respectively.

### Quantitative PCR for TERRAs

Gene expression changes were measured by qPCR using KAPA SYBR FAST One-Step Universal (cat# KK4651, Roche) according to the manufacturer’s instructions, with TERRA primer sequences sourced as per Vaid, *et al*.^56^ YWHAZ (*YWHAZ*) and GAPDH (*GAPDH*) were used as housekeeping genes (for all primer sequences, see **Supplementary Table 2**). Reverse transcription was performed at 45 °C for 5 min, enzyme activation at 95 °C for 3 min, denaturation at 95 °C for 3 s, annealing and extension at 60 °C for 20 s, for a total of 45 cycles. qPCR results were analysed using the 2^−ΔΔCT^ method^57^.

### Western blotting

Protein extraction and Western blotting were completed as previously described^58,59^. All antibody specifics can be found in **Supplementary Table 3**. Briefly, total protein was extracted from trophoblast organoids using radioimmunoprecipitation assay (RIPA) buffer (400 µL, ice-cold) supplemented with protease inhibitor cocktail (Roche) and phenylmethylsulfonyl fluoride (0.4 pmol). Organoids were incubated for 10 min at 4 °C, vortexed, and centrifuged (16,000 ×g, 10 min, 4 °C); supernatants were collected and protein concentration determined by Bradford assay (SpectraMax iD5, Molecular Devices, 595 nm).

Samples were denatured in Laemmli buffer with 2-mercaptoethanol (5 min, 95 °C) and separated on 4–20% Mini-PROTEAN TGX gels (Bio-Rad) alongside molecular weight markers. Electrophoresis was performed at 100 V for 1 h, and protein separation was confirmed by stain-free imaging (ChemiDoc Touch, Bio-Rad). Proteins were transferred to PVDF membranes overnight (27 V, 4 °C) in transfer buffer (25 mM Tris, 190 mM glycine, 20% methanol, pH 8.3).

Membranes were blocked in 5% skim milk/TBST (2 h, 25 °C), incubated with anti-NLRP3 polyclonal antibody (ab263899, Abcam; 1:300, 1 h, 25 °C), washed, and then probed with HRP-conjugated goat anti-rabbit IgG secondary antibody (Dako; 1:5000). After additional washes, membranes were developed with Clarity ECL substrate (#1705061, Bio-Rad, 5 min) and imaged using the ChemiDoc Touch system.

Band intensities were quantified with ImageLab software (Bio-Rad), normalized to β-actin protein, and corrected to an internal control (pooled trophoblast stem cells) on each membrane. Each sample was run in duplicate, and data were averaged across technical replicates. Quantification was performed from three independent experiments (n = 3 biological replicates).

### Immunofluorescence

Immunofluorescence was performed as previously described^2^, with some modifications. Briefly, organoids were rinsed with phosphate-buffered saline (PBS) and fixed using 4% paraformaldehyde in PBS for 20 minutes at room temperature. Permeabilization was performed using a solution containing 0.3% Triton X-100 and 3% BSA in PBS for 15 minutes, and blocking was carried out using a solution containing 3% BSA in PBS for 30 minutes. Then, organoids were incubated overnight with the primary antibody at 4 °C in PBS with 0.1% BSA at a dilution of 1:200 or 1:100. After three washes with PBS containing 0.1% Tween-20, organoids were incubated with secondary antibodies conjugated with Alexa Fluor 488 and Alexa Fluor 647 at a dilution of 1:500 for 30 minutes at room temperature. After three additional washes with the PBS-Tween-20 solution, nuclei were stained with NucBlue™ Fixed Cell ReadyProbes™ Reagent (DAPI) for 10 minutes, and the organoids were resuspended in 15 µL of Fluoromount-G mounting medium (#00-4958-02, Invitrogen). Organoids were imaged using an Olympus FV3000 confocal microscope, and images were analyzed with FV31S-SW FLUOVIEW software.

### gH2AX positivity

To calculate γH2AX positivity, immunofluorescence microscopy was performed as indicated above. Where γH2AX staining (red) was colocalised with cell nuclei (blue), this was considered a γH2AX-positive cell. For each sample, n=10 fields of view were imaged and the number of γH2AX-positive cells was counted. n=50 samples per condition.

### Immunohistochemistry and senescence assays

The ratio of syncytiotrophoblast to cytotrophoblast cells was assessed as an indicator of terminal differentiation and tissue aging. Undifferentiated cytotrophoblast cells are highly metabolically active in the term placenta whereas terminal differentiation into the syncytiotrophoblast indicates metabolic suppression^60^. Haemotoxylin and eosin immunohistochemical staining was employed to distinguish between cell types in term placenta sections of 10 μm. For each sample, n=10 fields of view were imaged and the number of syncytiotrophoblasts nuclei and cytotrophoblast nuclei were counted. n=50 samples per condition.

Cellular senescence was assessed using the senescence-associated β-Galactosidase (SA-β-Gal) Staining Kit (Abcam, ab65351), with small modifications. Namely, pan-cytokeratin was used to counterstain samples for trophoblast identification. Frozen placental sections or trophoblast organoids were air-dried at 37 °C for 30 min and fixed in SA-β-gal staining fixative for 15 min at room temperature. Samples were washed three times in PBS and incubated with SA-β-gal staining solution overnight at 37 °C. Following staining, samples were counterstained with pan-cytokeratin (Cytokeratin Pan Antibody Cocktail at 1:100 dilution, Thermo Fisher #MA5-13203) for 60 mins at 37 °C. Samples were then rinsed in running water (1 min), differentiated in 1% acid alcohol (10–20 s), and washed again under running water (1 min). Slides were dehydrated in graded alcohols, cleared in xylene, and mounted with coverslips.

Images were acquired by light microscopy at 40× magnification. SA-β-gal positivity was quantified using ImageJ software (NIH). The SA-β-gal–positive area was normalized to cytokeratin-positive total area. For each primary tissue sample (n=50 per group), 10 randomly selected regions were analysed, with quantification performed blinded to group allocation. For trophoblast organoids, n=50 whole organoids were analysed per group.

### Syncytial knot quantification

Trophoblast organoids were concurrently stained with DAPI (white; nucleus), Actin (cyan; cytoskeleton) and PSG1 (magenta; syncytiotrophoblast) (antibody details in **Supplementary Table 3**). Syncytial knots were defined as nuclear aggregates of at least 5 nuclei, as per Loukeris *et al*.^44^ Volumetric measurements (performed using ImageJ 3D suite as per Ollion, *et al.*^61^) yielded mean volumes of ∼445 µm³ for single syncytial nuclei and ∼2,200 µm³ for five-nuclei aggregates, consistent with previously reported estimates^62,63^. These findings highly agree with volumes based on previously mentioned diameters (11 µm) and elongation and flatness indices (0.6). All separate knot volumes were loaded into Microsoft Excel (Microsoft Corporation, Redmond, USA) and volumes smaller than 2400 µm^3^ were excluded to prevent inclusion of nuclear aggregates smaller than 5 nuclei. Syncytial knot density was subsequently calculated as the number of knots divided by the underlying cytotrophoblast volume. Total number of syncytial knots per organoid were counted, with n = 50 organoids analysed per group (control and LOPE). Quantification was performed blinded to experimental group.

### Angiogenic and inflammatory factor assays

Angiogenic factors sFlt-1 and PlGF were measured in culture media of trophoblast organoids as an indicator of preeclampsia pathology and severity. Factors were measured according to manufacturer’s instructions, using the PikoKine® ELISA kit for sFlt-1 (#MBS175839) and Human PlGF ELISA Kit from Quantikine (#DPG00), respectively.

Inflammatory factors IL-1β, IL-18, GSDMD and LDH were measured in maternal blood from control or LOPE pregnancies. Factors were measured according to manufacturer’s instructions. GSDMD was measured using the Human GSDMD ELISA Kit from Abcam (#ab272463). IL-1β, IL-18 and LDH were measured using the Human IL-1β ELISA Kit (#BMS224-2TEN), Human IL-18 ELISA Kit (#BMS267-2) and Lactate Dehydrogenase (LDH) Colorimetric Activity Kit (#EEA013), respectively; all from Invitrogen.

### Methylation array

All methylation array methods and results can be found in **Supplementary Files**.

### Statistics and reproducibility

All statistical analyses were performed using SPSS software and graphs were generated using GraphPad Prism. Data are presented as mean ± s.d. unless otherwise stated. For all experiments, statistical tests were two-sided and *p* < 0.05 was considered significant.

For telomere length assays, each measurement was conducted in technical triplicate, with trophoblast organoids generated from n = 3 biological samples per fetal sex, per treatment group (total n = 216), and repeated across four weeks of culture (final n = 1,080). For primary placental tissue, n = 50 per gestational week for both control and LOPE groups (total n = 600).

Quantitative PCR and Western blot data were obtained from three independent experiments (n = 3 biological replicates), each run in duplicate, and averaged across technical replicates. Senescence, oxidative stress, and DNA damage assays were quantified from n = 50 samples per condition and 10 fields of view per sample; syncytial knot quantification used n = 50 organoids per group, analysed blinded to condition.

Comparisons between two groups (e.g., control vs LOPE) were analysed by two-tailed Mann–Whitney tests. For multiple-group comparisons, Welch’s ANOVA with post hoc multiple-comparison correction was applied. Exact statistical tests and sample sizes are indicated in figure legends. No statistical method was used to predetermine sample size. All experiments were independently reproduced at least three times with consistent results.

## Authorship contributions

A.L.A. designed the study, performed experiments, collected and analysed data and wrote the manuscript. R.B. contributed to the experimental design and experimental work, collected data and edited the manuscript. M.D.S. performed bioinformatic analyses, wrote bioinformatics sections and reviewed the manuscript. D.Med. performed experimental work, collected data and reviewed the manuscript. E.M., G.M., J.M.W., L.H., J.M.P. and D.Mac. contributed to the experimental design and reviewed the manuscript. C.T.R. provided supervision, funding acquisition, review and editing of the manuscript. All authors contributed to the article and approved the submitted version.

## Funding

A.L.A. is supported by funding from Flinders University, the Flinders Foundation, the Channel 7 Children’s Research Foundation and a Future Making Fellowship from the University of Adelaide. C.T.R. is supported by an NHMRC Investigator Grant (GNT1174971) and a Matthew Flinders Fellowship from Flinders University. This work was supported by NIH NICHD R01 HD089685-01 Maternal molecular profiles reflect placental function and development across gestation PI Roberts.

**Supplementary Figure 1.**
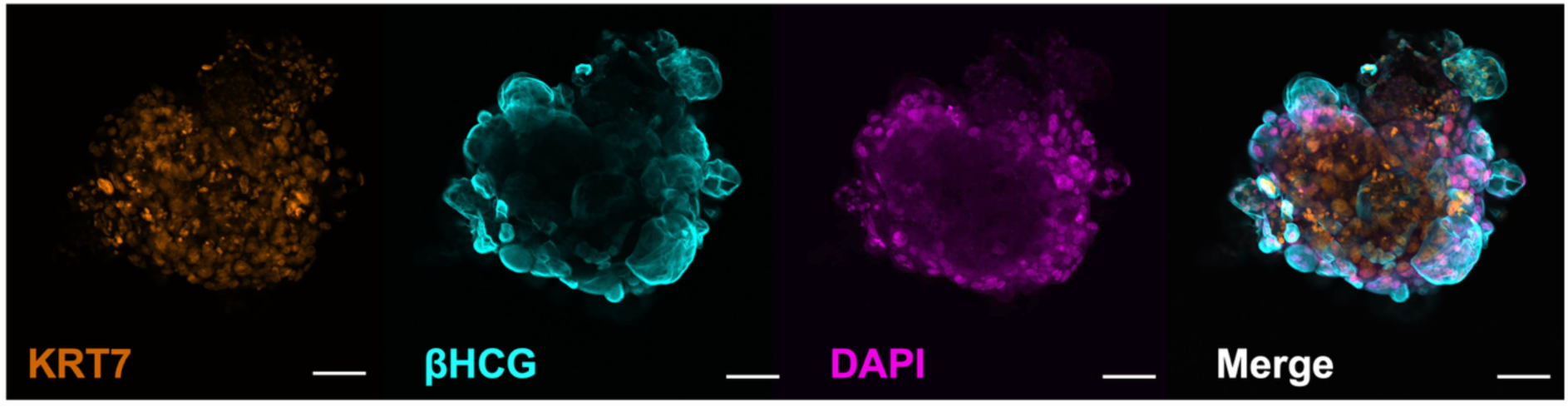
Trophoblast organoids were confirmed to exhibit correct polarity, with an outer layer of syncytiotrophoblast (βHCG, cyan) and underlying villous cytotrophoblasts (KRT7, orange). DAPI (magenta) marks cell nuclei. Scale bar = 100 μm.

**Supplementary Figure 2.**
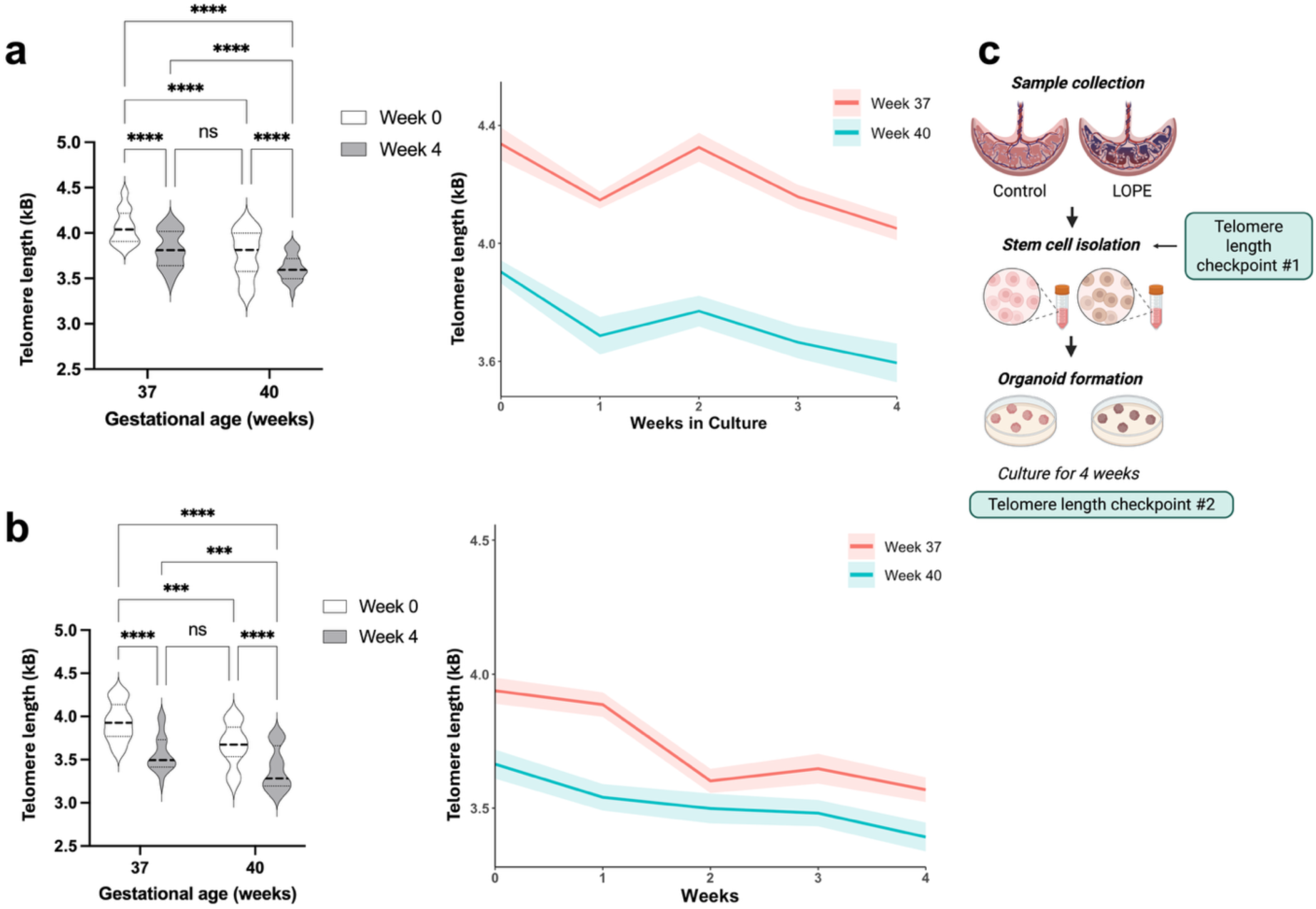
Telomere length was not significantly different in trophoblast organoids generated from 37 weeks’ gestation tissue after 4 weeks of culture compared with placenta tissue from 40 weeks’ gestation from **a**, control and **b**, LOPE participants. Data are presented as violin plots. **c**, experimental outline. **\*\*\*\*** p < 0.0001; ns = not significant.

**Supplementary Figure 3.**
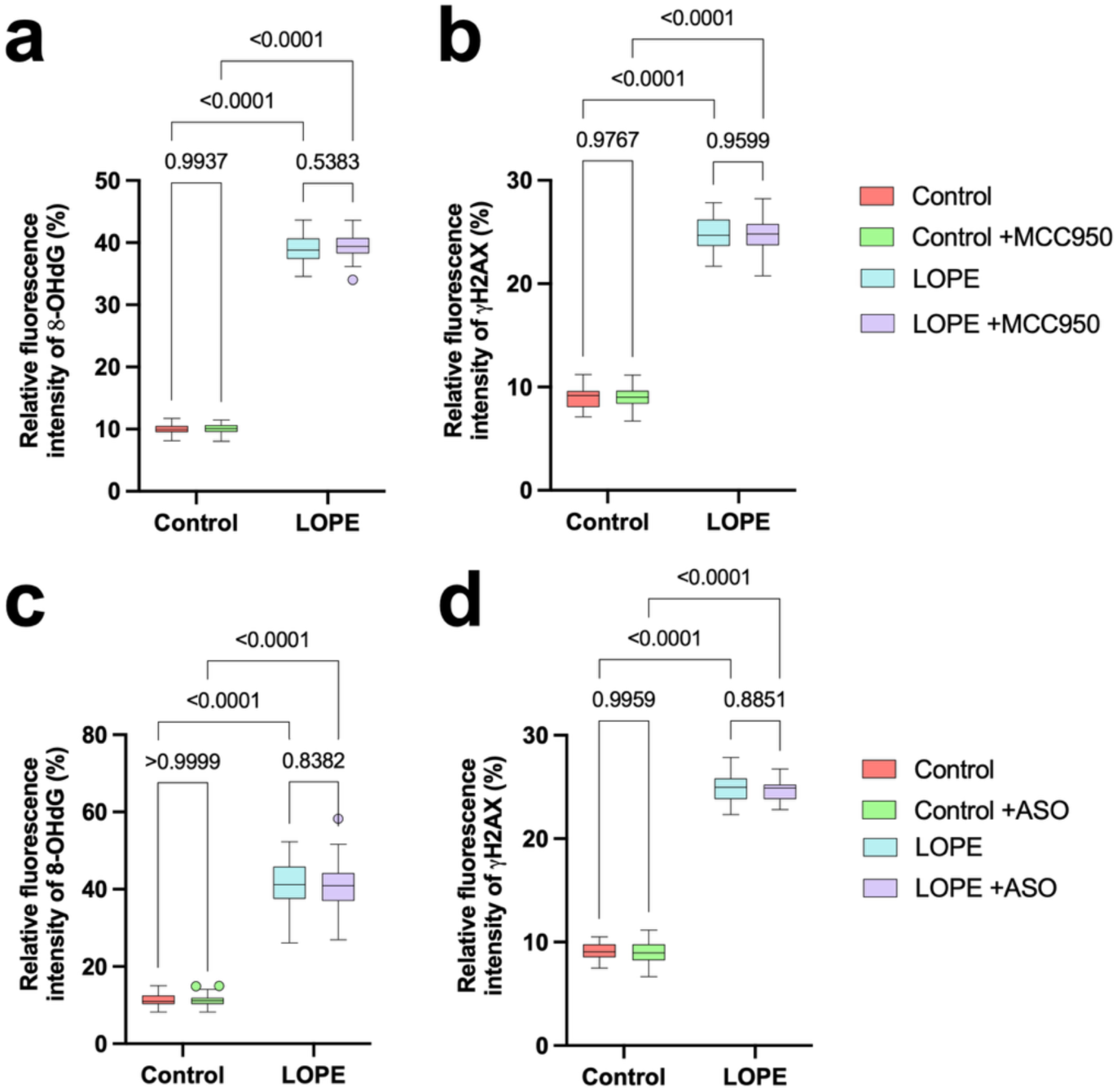
Relative fluorescence intensity for **a, c,** 8-OHdG and **b, d,** γH2AX was not significantly different in trophoblast organoids treated with **a, b,** MCC950 or **c, d,** ASOs targeting TERRAs; compared to untreated matched controls. Data presented as Tukey’s box and whisker plots.

**Supplementary Figure 4.**
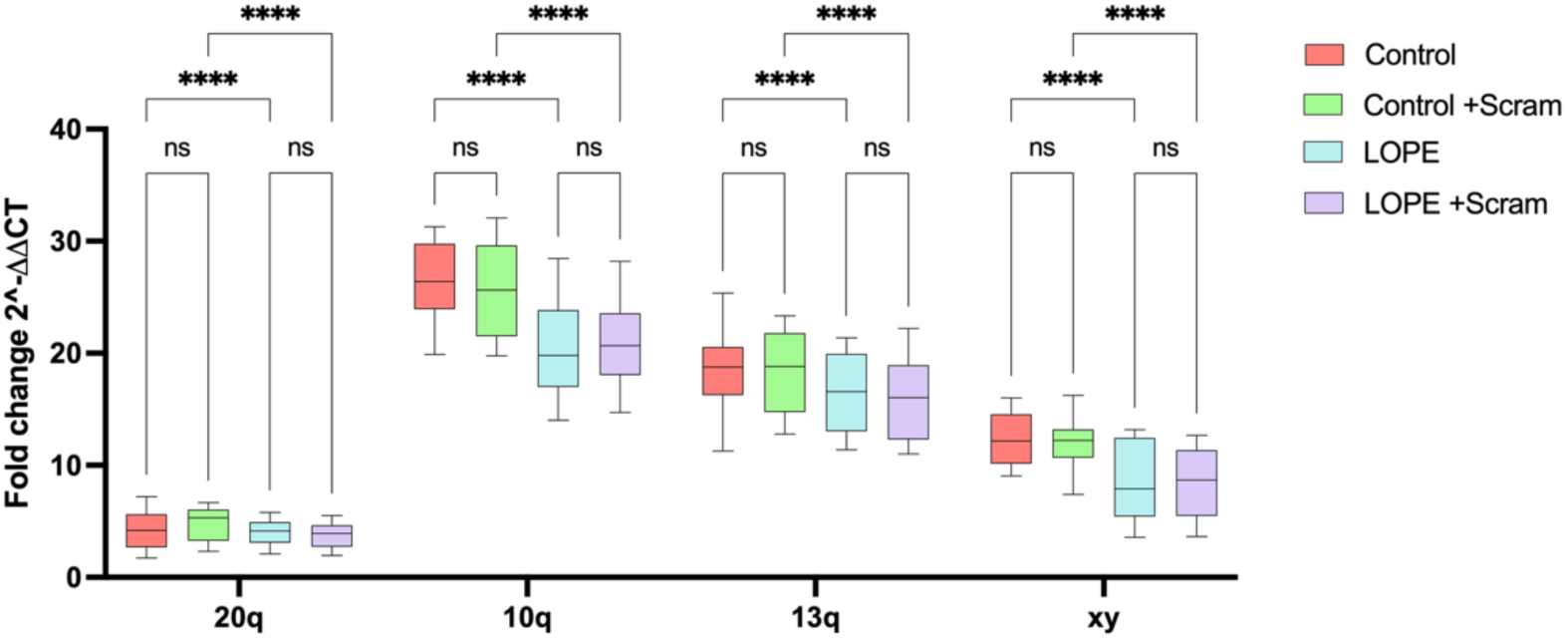
Expression of 20q, 10q, 13q and xy TERRAs was unchanged in both control and LOPE trophoblast organoids after treatment with the scrambled antisense oligonucleotide (ASO). Data presented as Tukey’s box and whisker plot.

**Supplementary Figure 5.**
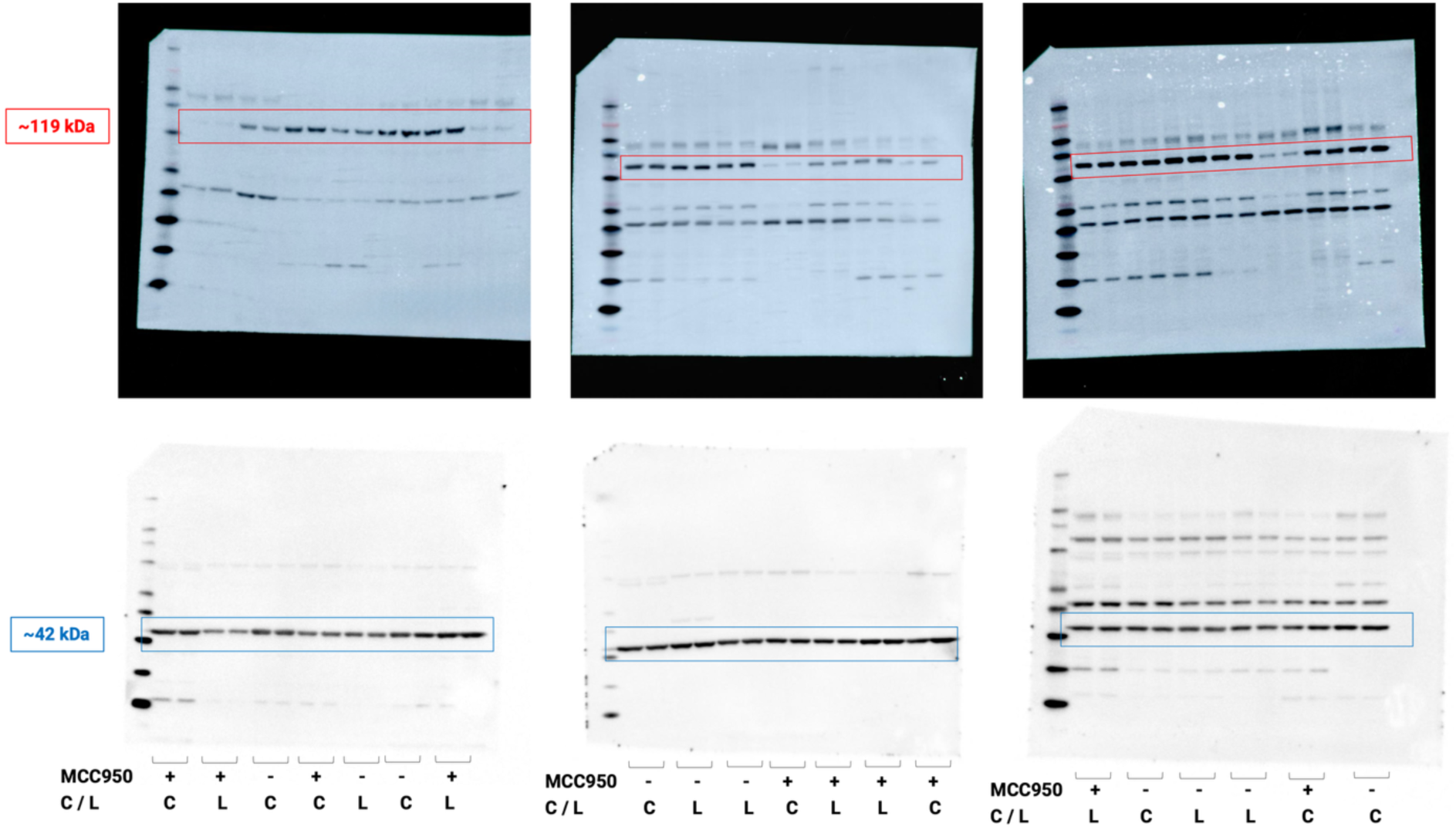
Uncropped Western Blots depicting NLRP3 protein (∼119 kDa) reduction with MCC950 treatment, and β-actin protein (∼42 kDa). C indicates control sample, L indicates LOPE sample.

**Supplementary Table 1.**
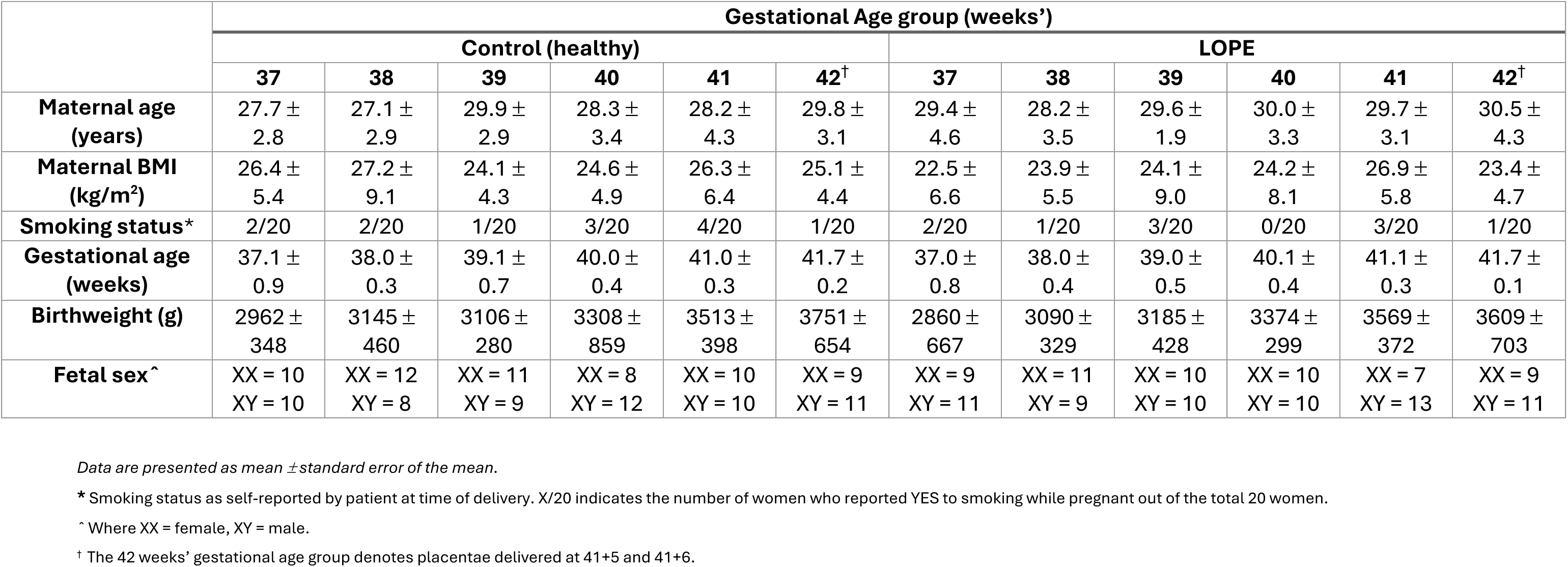
Characteristics of women who donated their placentae for this project. No significant differences found between parameters in control vs LOPE pregnancies.

|  | Gestational Age group (weeks') |  |  |  |  |  |  |  |  |  |  |  |
| --- | --- | --- | --- | --- | --- | --- | --- | --- | --- | --- | --- | --- |
|  | Control (healthy) |  |  |  |  |  | LOPE |  |  |  |  |  |
|  | 37 | 38 | 39 | 40 | 41 | 42<sup>†</sup> | 37 | 38 | 39 | 40 | 41 | 42<sup>†</sup> |
| **Maternal age (years)** | 27.7 ± 2.8 | 27.1 ± 2.9 | 29.9 ± 2.9 | 28.3 ± 3.4 | 28.2 ± 4.3 | 29.8 ± 3.1 | 29.4 ± 4.6 | 28.2 ± 3.5 | 29.6 ± 1.9 | 30.0 ± 3.3 | 29.7 ± 3.1 | 30.5 ± 4.3 |
| **Maternal BMI (kg/m<sup>2</sup>)** | 26.4 ± 5.4 | 27.2 ± 9.1 | 24.1 ± 4.3 | 24.6 ± 4.9 | 26.3 ± 6.4 | 25.1 ± 4.4 | 22.5 ± 6.6 | 23.9 ± 5.5 | 24.1 ± 9.0 | 24.2 ± 8.1 | 26.9 ± 5.8 | 23.4 ± 4.7 |
| **Smoking status\*** | 2/20 | 2/20 | 1/20 | 3/20 | 4/20 | 1/20 | 2/20 | 1/20 | 3/20 | 0/20 | 3/20 | 1/20 |
| **Gestational age (weeks)** | 37.1 ± 0.9 | 38.0 ± 0.3 | 39.1 ± 0.7 | 40.0 ± 0.4 | 41.0 ± 0.3 | 41.7 ± 0.2 | 37.0 ± 0.8 | 38.0 ± 0.4 | 39.0 ± 0.5 | 40.1 ± 0.4 | 41.1 ± 0.3 | 41.7 ± 0.1 |
| **Birthweight (g)** | 2962 ± 348 | 3145 ± 460 | 3106 ± 280 | 3308 ± 859 | 3513 ± 398 | 3751 ± 654 | 2860 ± 667 | 3090 ± 329 | 3185 ± 428 | 3374 ± 299 | 3569 ± 372 | 3609 ± 703 |
| **Fetal sex<sup>^</sup>** | XX = 10<br>XY = 10 | XX = 12<br>XY = 8 | XX = 11<br>XY = 9 | XX = 8<br>XY = 12 | XX = 10<br>XY = 10 | XX = 9<br>XY = 11 | XX = 9<br>XY = 11 | XX = 11<br>XY = 9 | XX = 10<br>XY = 10 | XX = 10<br>XY = 10 | XX = 7<br>XY = 13 | XX = 9<br>XY = 11 |

Data are presented as mean ± standard error of the mean.
\* Smoking status as self-reported by patient at time of delivery. X/20 indicates the number of women who reported YES to smoking while pregnant out of the total 20 women.
<sup>^</sup> Where XX = female, XY = male.
<sup>†</sup> The 42 weeks' gestational age group denotes placentae delivered at 41+5 and 41+6.

**Supplementary Table 2.**
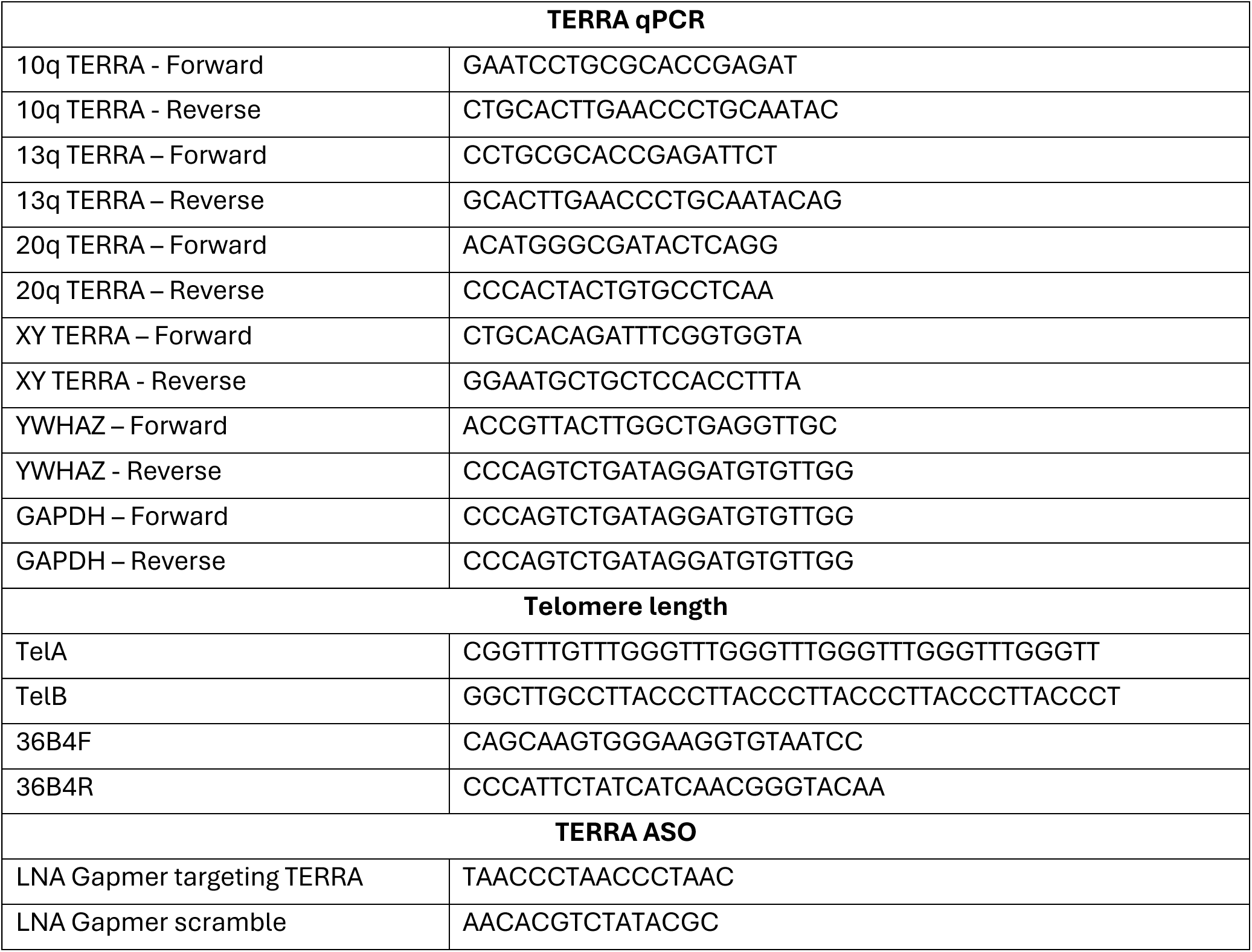
Sequences of oligos used for this project.

**Supplementary Table 3.** Catalogue numbers of antibodies used for this project.

| **Antibodies** | **Brand/Catalog** |
| --- | --- |
| Anti-hCGB Mouse Monoclonal Primary Antibody | Abcam (Ab9582) |
| Anti-Cytokeratin 7 Rabbit Polyclonal Antibody | Abcam (Ab181598) |
| Anti-PSG1 Rabbit Polyclonal Primary Antibody | Abcam (Ab233130) |
| Anti-gamma H2A.X (phospho S139) Mouse Monoclonal Primary Antibody | Abcam (Ab22551) |
| Anti-8-OHdG/8 Hydroxyguanosine Rabbit Polyclonal Primary Antibody | Bioss (Cat#BS-1278R) |
| Anti-Actin Mouse Monoclonal Antibody | Abcam (Ab11003) |
| Goat anti-Mouse IgG2a Cross-Adsorbed Secondary Antibody, Alexa Fluor™ 488 | ThermoFisher (Cat#A21131) |
| Goat anti-Rabbit IgG (H+L) Cross-Adsorbed Secondary Antibody, Alexa Fluor™ 488 | ThermoFisher (Cat#A11008) |
| Goat anti-Mouse IgG (H+L) Highly Cross-Adsorbed Secondary Antibody, Alexa Fluor™ Plus 647 | ThermoFisher (Cat#A32728) |

**Supplementary Table 4.** Characteristics of women who donated their placentae for methylation analyses stratified by fetal sex.

| Characteristic | Male (XY) | Female (XX) | Total |
| --- | --- | --- | --- |
| Sample size, n | 22 | 38 | 60 |
| Gestational age at delivery, weeks (mean ± SD) | 39.79 ± 1.37 | 39.84 ± 1.31 | 39.83 ± 1.32 |
| Delivery mode - LSCS in labour, n (%) | 6 (27.3) | 10 (26.3) | 16 (26.7) |
| Delivery mode - Operative vaginal, n (%) | 5 (22.7) | 5 (13.2) | 10 (16.7) |
| Delivery mode - Unassisted vaginal, n (%) | 9 (40.9) | 20 (52.6) | 29 (48.3) |
| Delivery mode - Prelabour LSCS, n (%) | 2 (9.1) | 3 (7.9) | 5 (8.3) |
| Preeclampsia, n (%) | 6 (27.3) | 10 (26.3) | 16 (26.7) |
| Maternal ethnicity* - Caucasian, n (%) | 21 (95.5) | 32 (84.2) | 53 (88.3) |
| Maternal ethnicity* - Asian, n (%) | 1 (4.5) | 1 (2.6) | 2 (3.3) |
| Maternal ethnicity* - Polynesian, n (%) | 0 (0) | 2 (5.3) | 2 (3.3) |
| Maternal ethnicity* - Aboriginal, n (%) | 0 (0) | 1 (2.6) | 1 (1.7) |
| Maternal ethnicity* - SouthEastAndFarEast, n (%) | 0 (0) | 2 (5.3) | 2 (3.3) |
| Maternal age, years (mean ± SD) | 23.82 ± 3.39 | 25.45 ± 4.52 | 24.85 ± 4.19 |
| Maternal BMI, kg/m<sup>2</sup> (mean ± SD) | 28.59 ± 7.54 | 27.01 ± 6.49 | 27.59 ± 6.88 |
| Birthweight, grams (mean ± SD) | 3621.86 ± 422.73 | 3449.18 ± 487.63 | 3512.5 ± 468.79 |
| Smoking during pregnancy<sup>†</sup>, n (%) | 1 (4.5) | 6 (15.8) | 7 (11.7) |

\* Ethnicity as self-reported by participants. This differs from the ethnicity used in the regression, which is predicted from the placental DNAm.
<sup>†</sup> Smoking status as self-reported by patient at time of delivery.

### Extended Data – Methylation analysis

#### Cell composition associations

We assessed whether cell composition was associated with fetal sex, gestational age and pregnancy outcome (late-onset preeclampsia versus uncomplicated). Using the ratio between two major trophoblast cell populations (cytotrophoblast (cyt) and syncytiotrophoblast (syn)). We first looked at the cyt:syn ratio between male and female fetal sex (**Supplementary Figure 6a**) and found no significant associations.

We further found no significant association between:

- total trophoblast (cytotrophoblasts and syncytiotrophoblasts combined) and fetal sex (**Supplementary Figure 6b**).
- combined endothelial and stromal cells and fetal sex (**Supplementary Figure 6c**), nor
- combined nucleated red blood cells and Hofbauer cells and fetal sex (**Supplementary Figure 6d**).

**Supplementary Figure 6.**
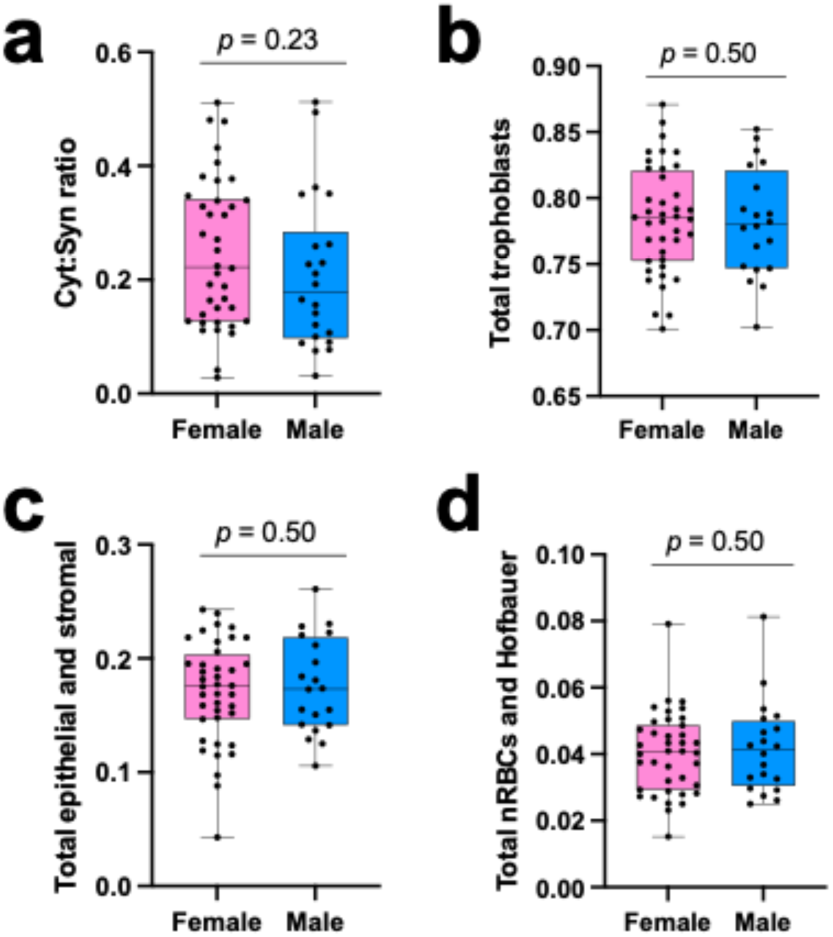
Sex-stratified associations of predicted **a,** cytotrophoblast to syncytiotrophoblast (cyt:syn) ratio, **b,** combined cytotrophoblast and syncytiotrophoblasts, **c,** combined endothelial and stromal cell proportions and **d,** combined nucleated red blood cells (nRBCs) and Hofbauer cells, all compared by fetal sex.

Previous literature has shown that the cyt:syn ratio will decrease as gestational age progresses^64^. To assess this in our own data, we calculated the Spearman’s correlation coefficient between cyt:syn ratio and clinically reported gestational age (**Supplementary Figure 7**). While there was a general downward trend in both male and female cyt:syn ratio, the negative correlation was only significant in females (R = ∼-0.35, *p* = 0.03). There was no statistically significant difference in clinically reported gestational age at birth between male and female (Mann-Whitney U test, *p* = 0.82).

**Supplementary Figure 7.**
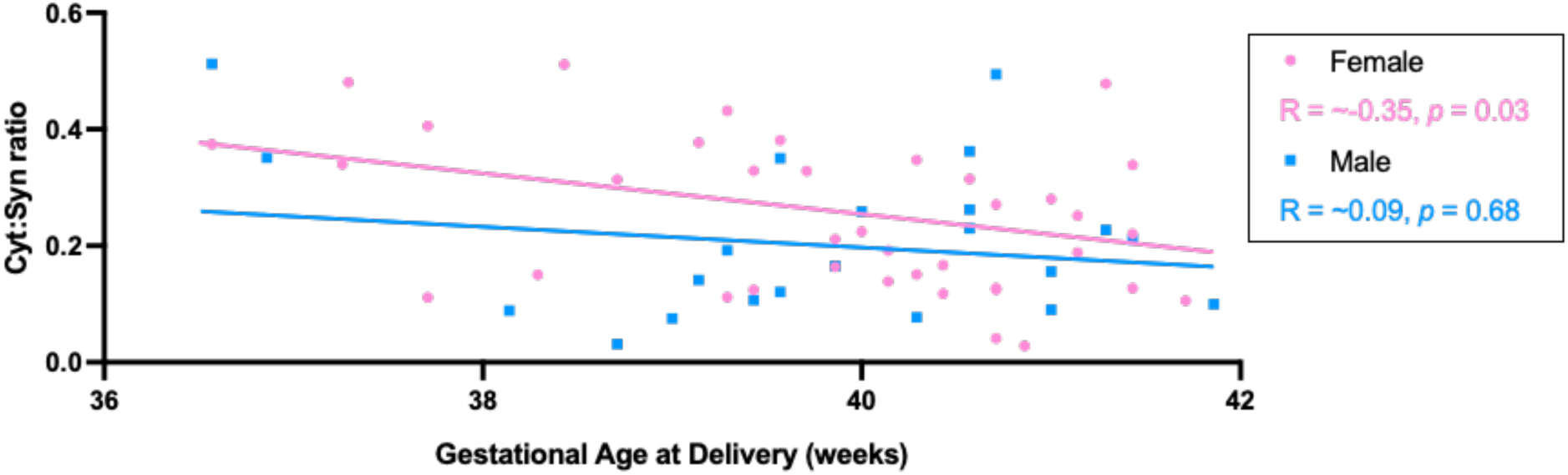
Association between placental cytotrophoblast-to-syncytiotrophoblast ratio and gestational age at delivery. PlaNET-predicted cell compositions were used to calculate the ratio of cytotrophoblast to syncytiotrophoblast cell content (cyt:syn) in bulk placental tissue samples. The scatter plot shows the relationship between cyt:syn ratio and gestational age at delivery for female (pink) and male (blue) placentae. Each point represents an individual sample, with solid lines indicating the fitted linear regression by fetal sex. A negative correlation between cyt:syn ratio and gestational age was observed in females (Spearman’s R = –0.35, p = 0.03), but not in males.

#### Epigenetic age acceleration associated with late onset preeclampsia

Late-onset preeclampsia has been associated with accelerated cellular aging in the human placenta^31,32^, prompting investigation into whether this extends to epigenetic aging. To explore this, we assessed epigenetic age acceleration using the Refined Robust Placental Clock (RRPC), calculating extrinsic epigenetic age acceleration (EEAA) as residuals from a linear regression of clinically reported gestational age on predicted epigenetic gestational age (epi-GA), and intrinsic epigenetic age acceleration (IEAA) with additional adjustment for PlaNET-predicted placental cell type proportions (Trophoblasts, Stromal, Hofbauer, Endothelial, nRBC, Syncytiotrophoblast). In the full cohort (n = 60), neither EEAA (**Supplementary Figure 8a**) nor IEAA (**Supplementary Figure 8c**) showed a significant association with preeclampsia outcome (EEAA: *p* = 0.77; IEAA: *p* = 0.65). Similarly, in sex-stratified analyses, no significant associations were observed. For EEAA (unadjusted for cell types; **Supplementary Figure 8b**), p-values were 0.56 for males and 0.43 for females. For IEAA (adjusted for cell types; **Supplementary Figure 8d**), p-values were 0.68 for males and 0.44 for females. These null findings suggest that, in this cohort, placental epigenetic age acceleration may not be a primary feature of late-onset preeclampsia, potentially due to sample size limitations or the specific disease pathology, warranting further investigation with larger datasets or alternative epigenetic markers.

**Supplementary Figure 8.**
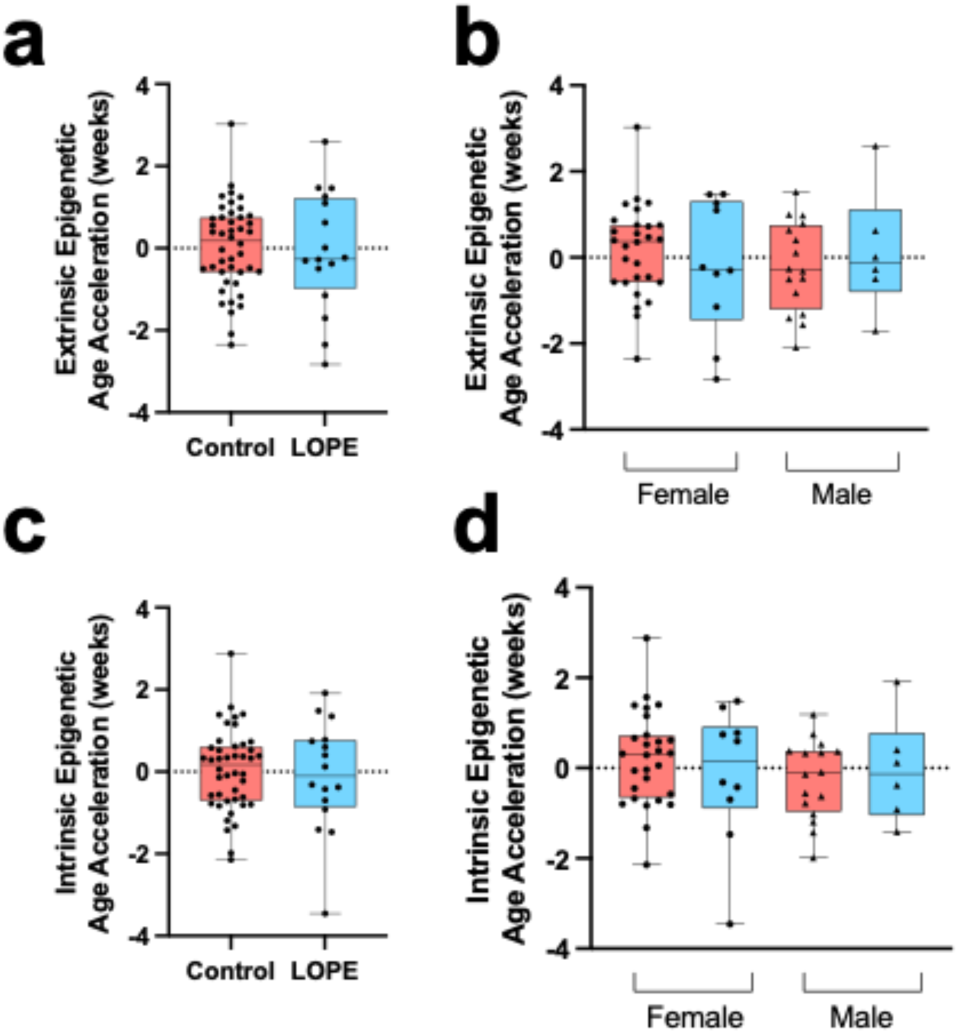
Extrinsic and intrinsic epigenetic age acceleration in placentae from late-onset preeclampsia (LOPE) and control pregnancies. Epigenetic gestational age (epi-GA) was estimated using the Refined Robust Placental Clock (RRPC), and epigenetic age acceleration was calculated as the residuals from a linear regression of clinically reported gestational age on epi-GA. **a,** Extrinsic epigenetic age acceleration (EEAA), unadjusted for cell type composition, compared between control (salmon) and LOPE (light blue) placentae. **b,** Sex-stratified EEAA showing comparisons within female and male placentae. **c,** Intrinsic epigenetic age acceleration (IEAA), adjusted for PlaNET-predicted placental cell type proportions (trophoblasts, stromal, Hofbauer, endothelial, nRBC, and syncytiotrophoblast), compared between control and LOPE placentae. **d,** Sex-stratified IEAA comparisons within female and male placentae. Boxplots display median and interquartile range, with individual data points overlaid. No significant differences were observed in EEAA or IEAA between LOPE and control groups in either the full cohort or sex-stratified analyses (all p > 0.4)

#### Differential methylation

We first assessed autosomal CpGs (n = 746,419) using a linear model adjusted for fetal sex, gestational age, and ancestry (Asian and Caucasian probabilities) to compare placental DNA methylation between late-onset preeclampsia and uncomplicated pregnancies (**Supplementary Figure 9**). No CpGs were differentially methylated (FDR < 0.05, |Δβ| > 0.05) between preeclamptic and uncomplicated pregnancies. Next, we considered all CpGs passing quality and filtering metrics (autosomal n = 746,419, ChrX = 16,769, ChrY = 318, total n = 763,506) using the same statistical thresholds and again found no significant CpGs. At relaxed FDR thresholds of 0.15 and 0.25, combined with |Δβ| > 0.03 (as used in other placental studies^65,66^), no CpGs were significant at FDR < 0.15. However, in both the autosomal and all chromosome models, at FDR < 0.25, a single CpG, cg23015112 (FDR = 0.16, Δβ = -0.06), was identified. This Chr15 probe is located in the body of the gene PLA2G4E, which encodes a phospholipase involved in lipid signalling. Further validation is warranted given the relaxed threshold. The absence of widespread differential methylation suggests that late-onset preeclampsia may not be strongly associated with global DNA methylation changes in this cohort, warranting exploration of region-specific analyses or alternative epigenetic mechanisms.

**Supplementary Figure 9.**
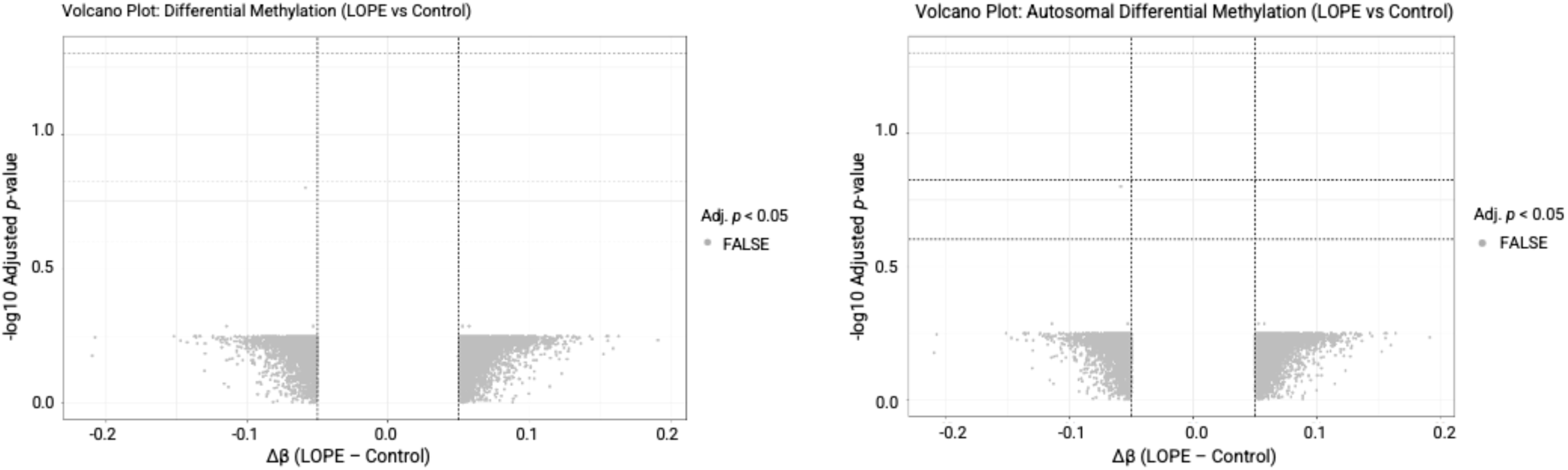
Volcano plots for differential DNAme in association with late-onset preeclampsia. More significant (lower FDR) values are shown at the top of the plot. Vertical dashed lines outline |Δβ| = 0.05, and horizontal dashed lines indicate FDR = 0.05, FDR = 0.15, and FDR = 0.25. [left] Volcano plot for all chromosomes. [right] Volcano plot for autosomes.

## Supplementary Methods – Methylation analysis

### Cohorts

60 placentae from two prospective cohorts were collected and processed for DNAme arrays in Adelaide, Australia. The two cohorts consisted of the SCOPE (SCreening fOr Pregnancy Endpoints) study^67^ and the Screening Tests to predict poor Outcomes of Pregnancy (STOP) study^48^ previously reported.

### Placental sampling

Placental sampling after delivery was conducted after assessment by the midwife to ensure tissue integrity. Briefly, the outer 2 cm perimeter was avoided, and tissue was dissected in a 2 cm-wide column extending inward from the edge. A 1 cm full-thickness block was taken at least 2 cm from both the placental edge and the cord insertion site, and four 0.5 cm³ pieces were dissected from the middle. Samples were immersed in RNA Later solution (Invitrogen) and frozen at -80°C for subsequent DNA extraction.

### DNAme arrays and data quality checks

DNA samples were purified and extracted using a modified version of the TES protocol (10mM Tris, 2mM Na_2_EDTA, 10% SDS)^68^ and hybridised to and processed on the Illumina Infinium Methylation EPIC BeadChip array according to the manufacturer’s protocol (Illumina, San Diego, CA, USA) at the PathWest Laboratory Medicine (QEII Medical Centre, Perth, Western Australia). Samples were distributed across array chips and rows with respect to pregnancy outcome (uncomplicated, late onset preeclampsia) and fetal sex variables to minimise potential batch effects.

### Epiphenotype estimation

DNAme data were read into R v 4.4.3 for processing. The PlaNET R package^69^ was used to determine DNAme-based estimates of gestational age at birth, genetic ancestry and placental cell type composition. These predicted estimates were calculated based on BMIQ-noob normalised data before probe filtering, as recommended in the PlaNET package documentation^69^ and as described by Khan *et al.*^64^ *Planet-predicted gestational age* can be calculated using 3 different built-in tools: The robust placental clock (RPC), the control placental clock (CPC) and the refined-robust placental clock (RRPC). We determined, given our samples were all collected at term and post-delivery, that the RRPC^70^, developed using samples from >36 weeks’ gestation and including pathological samples, was the appropriate measure for our analysis. *Planet-predicted genetic ancestry* is reported as a continuous variable on three axes of variation that sum to one, representing the contribution of African, East-Asian and European ancestry^69^. Planet-predicted cell-type composition was calculated using the robust partial corrections method, which yields compositional estimates of six major placental cell types (endothelial cells, stromal cells, Hofbauer cells, nucleated red blood cells, trophoblasts, and syncytiotrophoblasts)^71^. These methods follow those previously published by others^64^.

### Data processing

After epiphenotype variable estimation, the raw data were normalised for analysis using the adjustedDasen normalisation method^72^, chosen to avoid the introduction of sex bias into the autosomal data when normalising data from a mixed-sex cohort that includes sex chromosome data^72^. After normalisation, poor-quality probes (detection *p-*value > 0.01 or bead count < 3 or missing values in > 5% of samples) were removed from the dataset (*n*=2,993), as were cross-hybridising probes and probes (n=99,360)^73^. After data processing, a total of 763,506 CpGs (n=746,419 autosomal n=116,769 chrX; n=318 chrY) in 60 samples remained for downstream analysis.

### Statistical analysis

PlaNET predicted cell compositions were used to calculate the ratio of cytotrophoblast (PlaNET “trophoblasts”) to syncytiotrophoblast cell content (cyt:syn) in the bulk placental tissue samples. Cell type composition ratios were calculated as the proportion of cytotrophoblasts to syncytiotrophoblasts in bulk placental tissue samples. Two-sided Wilcoxon rank-sum tests were used to compare median cyt:syn ratios between fetal sex groups and between uncomplicated pregnancies versus preeclampsia cases. The association between cyt:syn ratios and gestational age at delivery was assessed using Spearman’s rank correlation coefficient.

The Refined Robust Placental Clock (RRPC) was used to estimate epigenetic gestational age, to assess whether late-onset preeclampsia was associated with epigenetic age acceleration. The clock was trained using samples from >36 weeks’ gestation and including pathological samples^70^. Using methods described in Beraldo, *et al.^C^*^5^ the gestational ages predicted by the RRPC clock are referred to herein as “epi-GA”. Similarly, *Extrinsic epigenetic age acceleration* was calculated as the residuals of a linear regression model with clinically reported gestational age at birth as the independent variable and epi-GA as the dependent variable to provide an overall picture of epigenetic aging. *Intrinsic epigenetic age acceleration* was calculated in the same manner, but was also adjusted for PlaNET-predicted cell composition effects. The association between epigenetic age acceleration and late-onset preeclampsia was evaluated using linear models, both in the whole cohort and sex-stratified analyses.

DNAme data (β values) at all filtered autosomal CpGs (n = 763,506) were converted to M values prior to linear modelling to test for DNAme differences between preeclamptic and uncomplicated placentae. Fetal sex, gestational age at delivery, and two of the three PlaNET ancestry estimates were included as additive covariates (probability of Asian ancestry and probability of European ancestry were selected for inclusion, given that the probability of African ancestry estimates were extremely low in these cohorts and that the predicted ancestry matched the self-reported ancestry perfectly). Clinical gestational age (calculated from last menstrual period) was used in all statistical models.

Linear models were run using the *limma* R package^74^. We applied a delta beta (|Δβ|) cutoff of ≥0.05 to identify biologically meaningful methylation differences, consistent with commonly used thresholds in epigenome-wide association studies^75,76^. This threshold represents a 5% difference in methylation levels and helps distinguish biologically relevant changes from technical noise in array-based methylation studies.

## References

1 Harman, D. Aging: overview. Annals of the New York Academy of Sciences 928, 1–21 (2001).

2 Arthurs, A. L. et al. Circular RNAs accumulate in aging human placental tissue and in stillbirth, leading to DNA damage and cellular senescence. American Journal of Obstetrics & Gynecology (2025). 10.1016/j.ajog.2025.08.030

3 Chen, Z. et al. Advanced maternal age causes premature placental senescence and malformation via dysregulated α-Klotho expression in trophoblasts. Aging Cell 20, e13417 (2021).

4 Bozack, A. K. et al. DNA methylation age at birth and childhood: performance of epigenetic clocks and characteristics associated with epigenetic age acceleration in the Project Viva cohort. Clinical epigenetics 15, 62 (2023).

5 Huff, K. K., Roell, K. R., Eaves, L. A., O’Shea, T. M. & Fry, R. C. Prenatal Exposure to Metals Is Associated with Placental Decelerated Epigenetic Gestational Age in a Sex-Dependent Manner in Infants Born Extremely Preterm. Cells 14, 306 (2025).

6 Habtewold, T. D. et al. Longitudinal maternal glycemia during pregnancy and placental epigenetic age acceleration. Clinical Epigenetics 17, 19 (2025).

7 Workalemahu, T., Shrestha, D., Tajuddin, S. M. & Tekola-Ayele, F. Maternal cardiometabolic factors and genetic ancestry influence epigenetic aging of the placenta. Journal of developmental origins of health and disease 12, 34–41 (2021).

8 Saeed, H., Wu, J., Tesfaye, M., Grantz, K. L. & Tekola-Ayele, F. Placental accelerated aging in antenatal depression. American journal of obstetrics & gynecology MFM 6, 101237 (2024).

9 Saenen, N. et al. Air pollution-induced placental alterations: an interplay of oxidative stress, epigenetics, and the aging phenotype? Clinical epigenetics 11, 124 (2019).

10 Brosens, I., Pijnenborg, R., Vercruysse, L. & Romero, R. The “Great Obstetrical Syndromes” are associated with disorders of deep placentation. American journal of obstetrics and gynecology 204, 193–201 (2011).

11 Burton, G. J., Redman, C. W., Roberts, J. M. C Moffett, A. Pre-eclampsia: pathophysiology and clinical implications. Bmj 366 (2019).

12. Audette, M. C. & Kingdom, J. C. in Seminars in Fetal and Neonatal Medicine. 119-125 (Elsevier).

13 Kim, Y. M. et al. Failure of physiologic transformation of the spiral arteries in patients with preterm labor and intact membranes. American journal of obstetrics and gynecology 189, 1063–1069 (2003).

14 Maiti, K. et al. Evidence that fetal death is associated with placental aging. Am J Obstet Gynecol 217, 441 (2017).

15 Neiger, R. Long-term effects of pregnancy complications on maternal health: a review. Journal of clinical medicine 6, 76 (2017).

16 Smith, G. C., Pell, J. P. & Walsh, D. Pregnancy complications and maternal risk of ischaemic heart disease: a retrospective cohort study of 129 290 births. The Lancet 357, 2002–2006 (2001).

17 Magnussen, E. B. et al. Prepregnancy cardiovascular risk factors as predictors of pre-eclampsia: population based cohort study. BMJ 335, 978 (2007). 10.1136/bmj.39366.416817.BE

18 Bellamy, L., Casas, J.-P., Hingorani, A. D. & Williams, D. Type 2 diabetes mellitus after gestational diabetes: a systematic review and meta-analysis. The lancet 373, 1773–1779 (2009).

19 Dipla, K. et al. Impairments in microvascular function and skeletal muscle oxygenation in women with gestational diabetes mellitus: links to cardiovascular disease risk factors. Diabetologia 60, 192–201 (2017).

20 Smith, G. D., Harding, S. & Rosato, M. Relation between infants’ birth weight and mothers’ mortality: prospective observational study. Bmj 320, 839–840 (2000).

21 Kim, M. K. et al. Socioeconomic status can affect pregnancy outcomes and complications, even with a universal healthcare system. International journal for equity in health 17, 2 (2018).

22 Arthurs, A. L., Harrison, J. K., Williamson, J. M. & Roberts, C. T. Pregnancy and Birth Trends Across Australia, the United States of America and the United Kingdom. Journal of Clinical Medicine 14, 5841 (2025).

23 Peacock, J. L., Bland, J. M. & Anderson, H. R. Preterm delivery: effects of socioeconomic factors, psychological stress, smoking, alcohol, and caffeine. Bmj 311, 531–535 (1995).

24 Silva, L. M. et al. Low socioeconomic status is a risk factor for preeclampsia: the Generation R Study. Journal of hypertension 26, 1200–1208 (2008).

25 Magee, L. A. et al. The 2021 International Society for the Study of Hypertension in Pregnancy classification, diagnosis & management recommendations for international practice. Pregnancy hypertension 27, 148–169 (2022).

26 Obstetricians, A. C. o. & Gynecologists. Hypertension in pregnancy. Report of the American College of Obstetricians and Gynecologists’ task force on hypertension in pregnancy. Obstet. Gynecol. 122, 1122 (2013).

27 Lisonkova, S. & Joseph, K. Incidence of preeclampsia: risk factors and outcomes associated with early-versus late-onset disease. American journal of obstetrics and gynecology 209, 544. e541–544. e512 (2013).

28 Jauniaux, E., Hempstock, J., Greenwold, N. & Burton, G. J. Trophoblastic oxidative stress in relation to temporal and regional differences in maternal placental blood flow in normal and abnormal early pregnancies. The American journal of pathology 162, 115–125 (2003).

29 Myatt, L. & Webster, R. P. Vascular biology of preeclampsia. Journal of Thrombosis and Haemostasis 7, 375–384 (2009).

30 Sheridan, R. M., Stanek, J., Khoury, J. & Handwerger, S. Abnormal expression of transcription factor activator protein-2α in pathologic placentas. Human pathology 43, 1866–1874 (2012).

31 Farladansky-Gershnabel, S. et al. 317: Senescence and telomere shortening are enhanced in placentas from pregnancies with early compared to late onset preeclampsia. American Journal of Obstetrics & Gynecology 218, S199 (2018).

32 Ortega, M. A. et al. Placental Tissue Calcification and Its Molecular Pathways in Female Patients with Late-Onset Preeclampsia. Biomolecules 14, 1237 (2024).

33 Sultana, Z., Maiti, K., Dedman, L. & Smith, R. Is there a role for placental senescence in the genesis of obstetric complications and fetal growth restriction? American journal of obstetrics and gynecology 218, S762–S773 (2018).

34 Christiansen, L. et al. DNA methylation age is associated with mortality in a longitudinal Danish twin study. Aging cell 15, 149–154 (2016).

35 Bhatti, G. et al. Placental epigenetic clocks derived from crowdsourcing: Implications for the study of accelerated aging in obstetrics. Iscience 28 (2025).

36 Mayne BT, L. S., Smith AK, Breen J, Roberts CT, Bianco-Miotto T. Accelerated placental aging in early onset preeclampsia pregnancies identified by DNA methylation. Epigenomics 9, 279–289 (2017). doi:10.2217/epi-2016-0103

37 Cruz, J. d. O., Conceicao, I. M., Tosatti, J. A., Gomes, K. B. & Luizon, M. R. Global DNA methylation in placental tissues from pregnant with preeclampsia: a systematic review and pathway analysis. Placenta 101, 97–107 (2020).

38 Raijmakers, M. T., Dechend, R. & Poston, L. Oxidative stress and preeclampsia: rationale for antioxidant clinical trials. Hypertension 44, 374–380 (2004).

39 Zeisler, H. et al. Predictive value of the sFlt-1: PlGF ratio in women with suspected preeclampsia. N Engl J Med 374, 13–22 (2016).

40 Maynard, S., Maynard SE, Min JY, Merchan J, Lim KH, Li J, et al. Excess placental soluble fms-like tyrosine kinase 1 (sFlt1) may contribute to endothelial dysfunction, hyperten-sion, and proteinuria in preeclampsia. J Clin Invest 111, 649–658 (2003).

41 Weel, I. C. et al. Increased expression of NLRP3 inflammasome in placentas from pregnant women with severe preeclampsia. Journal of reproductive immunology 123, 40–47 (2017).

42 Xu, L. et al. The NLRP3 rs10754558 polymorphism is a risk factor for preeclampsia in a Chinese Han population. The Journal of Maternal-Fetal & Neonatal Medicine 32, 1792–1799 (2019).

43 Pontillo, A. et al. NLRP1 L155H polymorphism is a risk factor for preeclampsia development. American Journal of Reproductive Immunology 73 (2015).

44 Loukeris, K., Sela, R. & Baergen, R. N. Syncytial knots as a reflection of placental maturity: reference values for 20 to 40 weeks’ gestational age. Pediatric and developmental pathology 13, 305–309 (2010).

45 Menkhorst, E. et al. IL11 activates the placental inflammasome to drive preeclampsia. Frontiers in Immunology 14, 1175926 (2023).

46 Cheng, S.-B. et al. Pyroptosis is a critical inflammatory pathway in the placenta from early onset preeclampsia and in human trophoblasts exposed to hypoxia and endoplasmic reticulum stressors. Cell death & disease 10, 927 (2019).

47 Deng, Z. et al. Formation of telomeric repeat-containing RNA (TERRA) foci in highly proliferating mouse cerebellar neuronal progenitors and medulloblastoma. Journal of cell science 125, 4383–4394 (2012).

48 Mohammed, H. et al. Safety and protective effects of maternal influenza vaccination on pregnancy and birth outcomes: A prospective cohort study. EClinicalMedicine 26, 100522 (2020).

49 Yang, L. et al. Innate immune signaling in trophoblast and decidua organoids defines differential antiviral defenses at the maternal-fetal interface. Elife 11, e79794 (2022).

50 Perera, A. P. et al. MCC950, a specific small molecule inhibitor of NLRP3 inflammasome attenuates colonic inflammation in spontaneous colitis mice. Scientific reports 8, 8618 (2018).

51 Bettin, N. et al. TERRA transcripts localize at long telomeres to regulate telomerase access to chromosome ends. Science Advances 10, eadk4387 (2024).

52 Chu, H.-P. et al. TERRA RNA antagonizes ATRX and protects telomeres. Cell 170, 86–101. e116 (2017).

53 Tominaga, T. et al. Exogenously-added copper/zinc superoxide dismutase rescues damage of endothelial cells from lethal irradiation. Journal of Clinical Biochemistry and Nutrition 50, 78–83 (2011).

54 Arai, Y. et al. Inflammation, but not telomere length, predicts successful ageing at extreme old age: a longitudinal study of semi-supercentenarians. EBioMedicine 2, 1549–1558 (2015).

55 Martin, N. A. et al. Monochrome multiplex quantitative PCR telomere length measurement. Journal of visualized experiments: JoVE, 10.3791/66545 (2024).

56 Vaid, R. et al. METTL3 drives telomere targeting of TERRA lncRNA through m6A-dependent R-loop formation: a therapeutic target for ALT-positive neuroblastoma. Nucleic Acids Research 52, 2648–2671 (2024).

57 Livak, K. J. & Schmittgen, T. D. Analysis of relative gene expression data using real-time quantitative PCR and the 2− ΔΔCT method. methods 25, 402–408 (2001).

58 Arthurs, A., Lumbers ER, Delforce SJ, Mathe A, Morris BJ, Pringle KG. The role of oxygen in the regulation of miRNAs in control of the placental renin-angiotensin system. Molecular Human Reproduction 25, 206–217 (2019).

59 Arthurs, A. L. et al. Genetically edited human placental organoids cast new light on the role of ACE2. Cell Death & Disease 16, 78 (2025).

60 Kolahi, K. S., Valent, A. M. & Thornburg, K. L. Cytotrophoblast, not syncytiotrophoblast, dominates glycolysis and oxidative phosphorylation in human term placenta. Scientific reports 7, 42941 (2017).

61 Ollion, J., Cochennec, J., Loll, F., Escudé, C. & Boudier, T. TANGO: a generic tool for high-throughput 3D image analysis for studying nuclear organization. Bioinformatics 29, 1840–1841 (2013).

62 Bierens, J. et al. Imaging and quantification of placental terminal villi microvasculature and nuclear characteristics in preeclampsia. European Journal of Obstetrics & Gynecology and Reproductive Biology 308, 181–189 (2025).

63 Chua, S., Wilkins, T., Sargent, I. C Redman, C. Trophoblast deportation in pre-eclamptic pregnancy. BJOG: An International Journal of Obstetrics & Gynaecology 98, 973–979 (1991).

64 Khan, A. et al. The application of epiphenotyping approaches to DNA methylation array studies of the human placenta. Epigenetics & Chromatin 16, 37 (2023).

65 Beraldo, E. O. et al. Exposure to prenatal maternal stress is associated with epigenetic age acceleration and altered cell composition in the placenta: The QF2011 Queensland Flood Study. Placenta (2025).

66 Inkster, A. M. et al. Profiling placental DNA methylation associated with maternal SSRI treatment during pregnancy. Scientific Reports 12, 22576 (2022).

67 North, R. A. et al. Clinical risk prediction for pre-eclampsia in nulliparous women: development of model in international prospective cohort. Bmj 342 (2011).

68 Miller, S. A., Dykes, D. D. & Polesky, H. A simple salting out procedure for extracting DNA from human nucleated cells. Nucleic acids research 16, 1215 (1988).

69 Yuan, V. et al. Accurate ethnicity prediction from placental DNA methylation data. Epigenetics & chromatin 12, 51 (2019).

70 Lee, Y. et al. Placental epigenetic clocks: estimating gestational age using placental DNA methylation levels. Aging (Albany NY) 11, 4238 (2019).

71 Yuan, V. et al. Cell-specific characterization of the placental methylome. BMC genomics 22, 6 (2021).

72 Wang, Y. et al. InterpolatedXY: a two-step strategy to normalize DNA methylation microarray data avoiding sex bias. Bioinformatics 38, 3950–3957 (2022).

73 Zhou, W., Laird, P. W. & Shen, H. Comprehensive characterization, annotation and innovative use of Infinium DNA methylation BeadChip probes. Nucleic acids research 45, e22–e22 (2017).

74 Ritchie, M. E. et al. limma powers differential expression analyses for RNA-sequencing and microarray studies. Nucleic acids research 43, e47–e47 (2015).

75 Du, P. et al. Comparison of Beta-value and M-value methods for quantifying methylation levels by microarray analysis. BMC bioinformatics 11, 587 (2010).

76 Tsai, P.-C. & Bell, J. T. Power and sample size estimation for epigenome-wide association scans to detect differential DNA methylation. Int J Epidemiol 44, 1429–1441 (2015).

